# An Asgard archaeon from a modern analog of ancient microbial mats

**DOI:** 10.1101/2025.07.22.663070

**Authors:** Stephanie-Jane Nobs, Matthew D. Johnson, Timothy J. Williams, Julia Meltzer, Xabier Vázquez-Campos, Fraser I. MacLeod, Keiran Rowell, Miranda Pitt, Bindusmita Paul, Doulin C. Shepherd, Katharine A. Michie, Iain G. Duggin, Debnath Ghosal, Brendan P. Burns

## Abstract

It has been proposed that eukaryotic cells evolved via symbiosis between sulfate-reducing bacteria and hydrogen-producing archaea. Here we describe a highly enriched culture of a novel Asgard archaeon, *Nerearchaeum marumarumayae*, with a bacterium *Stromatodesulfovibrio nilemahensis* from a stromatolite-associated microbial mat. The *N. marumarumayae* genome indicates it produces H_2_, acetate, formate, and sulfite, while *S. nilemahensis* synthesizes amino acids and vitamins, which can be exchanged in a syntrophic partnership. Electron cryotomography revealed *N. marumarumayae* cells produce chains of budded envelope vesicles attached to the coccoid cell body by extracellular fibers, and intracellular tube- and cage-like structures. Furthermore, the two species were observed interacting via intercellular nanotubes assembled by the bacterium. These characteristics and interactions may reflect an early step in the symbiotic evolution of eukaryotic cells.

## Introduction

A remaining mystery in biology is understanding how the first eukaryotic cells evolved. Eukaryogenesis is thought to have occurred ∼2 billion years (Ga) ago from a symbiotic merger of an ancestral archaeal cell and a bacterium, but the nature of their interactions and transitions to complete interdependence are unknown (*1*). Growing evidence suggests that the archaeal progenitor is related to recently discovered organisms known as Asgard archaea, which are considered the closest extant relatives of eukaryotes (*2, 3*). Asgard archaea encode a plethora of proteins involved in classically eukaryotic functions, such as endomembrane trafficking and associated intracellular signaling. Our understanding of the biology of Asgard archaea and their implications for eukaryogenesis have been limited by their relative scarcity, slow growth, and difficulty of cultivation (*4-6*).

It has been proposed that eukaryogenesis evolved in complex communities of microbial mats—thick layered biofilms in which different organisms may interact and exchange compounds to acquire energy and nutrients, developing metabolic interdependencies essential for survival (syntrophy) (*7*). Microbial mats were widespread in the Proterozoic Eon (2.5-0.5 Ga ago) (*8*) and persist today in environments where eukaryotic grazing is limited, such as the hypersaline (∼7% salt) (*9*) intertidal mats of Shark Bay (*Gathaagudu*) in Western Australia (Fig. 1A). These mats are dynamic stratified environments with oxygenic upper layers dominated by cyanobacteria, and anaerobic bacteria and archaea in the subsurface, including several Asgard lineages referred to as Loki-, Thor-, Odin- and Heimdallarchaeota (*10, 11*).

**Figure 1.**
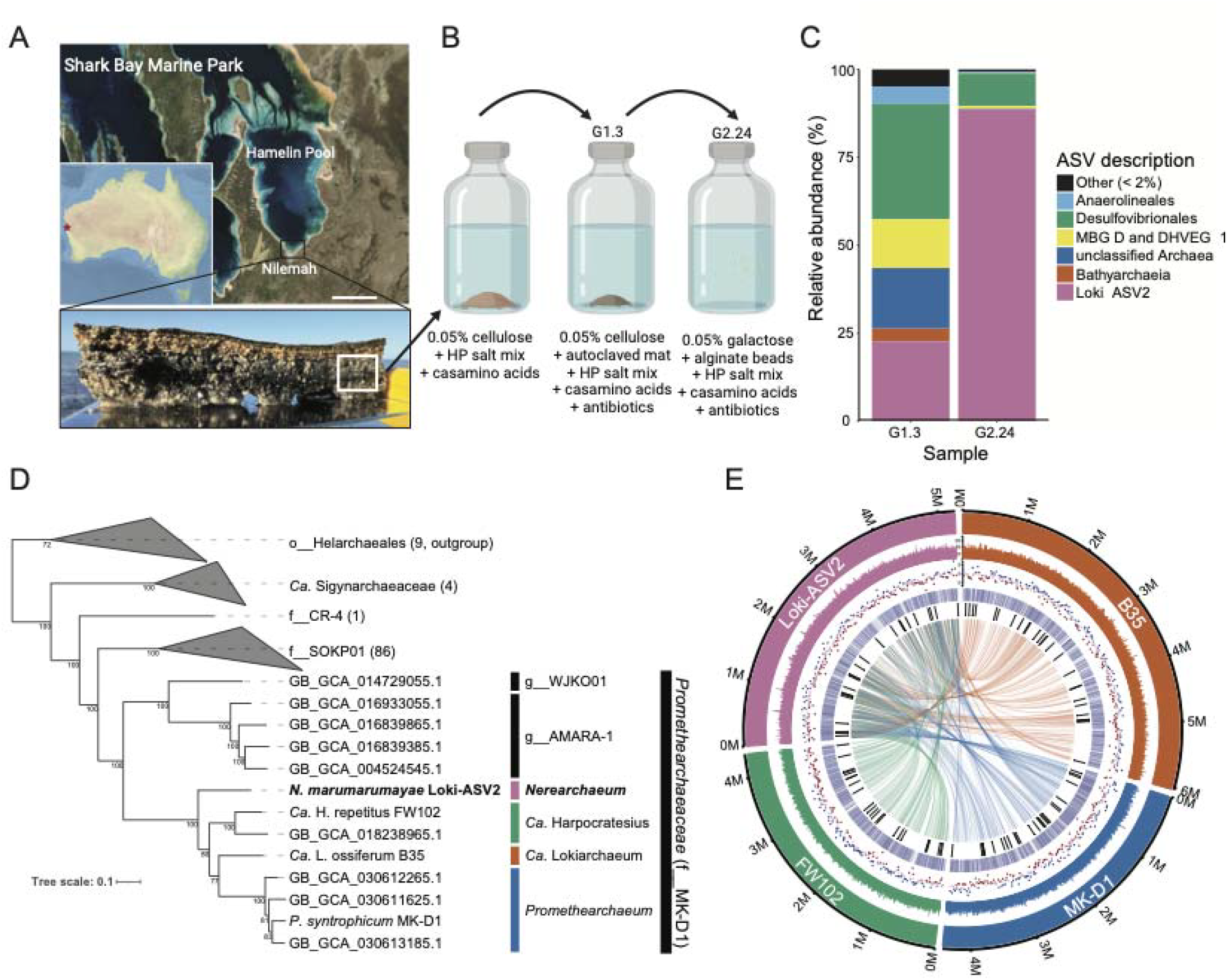
Enrichment of an Asgard archaeon from microbial mats. **(A)** Location map and satellite image of Shark Bay, Australia, indicating the sampling location Nilemah. Scale bar: 20 km (Source: Geoscience Australia). The lower photograph shows a smooth mat sample in cross-section with visible layering and the approximate anoxic region (white box) used to start cultures. **(B)** Schematic of culturing pipeline (made with BioRender). **(C)** Composition of representative cultures including one with the highest enrichment (G2.24) of a single Asgard archaeal strain (Loki-ASV2) as determined by 16S rRNA gene amplicon sequencing. **(D)** Phylogenic placement of *Nerearchaeum marumarumayae* Loki-ASV2 in a maximum-likelihood tree based on a concatenated set of marker protein sequences compared to the indicated Asgard archaea. Species, MAGs, or groups are labeled according t GTDB names and taxon level letter tags. **(E)** CIRCOS plot comparing *N. marumarumayae* Loki-ASV2 with the indicated *Promethearchaeaceae* circular genomes. The tracks indicate, from outmost to the inside: GC content (%), GC bias (blue: positive; red: negative), orthologous proteins, genes encoding ESP, and shared syntenic blocks (colors indicate the genome sharing with Loki-ASV2).

### The novel Asgard *N. marumarumayae* enriched to high abundance from microbial mats

To begin characterizing Asgard archaea from Hamelin Pool in Shark Bay, mat subsurface samples (Fig. 1A) were used as culture inoculum in anaerobic media containing Hamelin Pool (HP) salts, supplemented with various carbohydrates and amino acids, trace nutrients, and antibiotics (Fig. 1B, materials and methods) to support Asgard archaeal growth and inhibit bacterial growth (*4, 5*). After incubation of these cultures and subsequent subcultures (at 30 °C for three months each), flocs that were readily resuspended were often observed. Amplicon sequencing of the 16S rRNA gene in these cultures revealed enrichments of up to 31% of all detected amplicon sequencing variants (ASVs) from the *Promethearchaeaceae* family (f__MK-D1) of Asgard archaea. In one culture (G1.3), we detected only one *Promethearchaeaceae* member, which we designated Loki-ASV2 in reference to its similarity to the original Asgard sequences from Loki’s Castle hydrothermal vents (*12*) (Fig. 1C).

Additional subcultures from G1.3 exhibited further enrichment, including G2.24 with 89% Loki-ASV2 after six months incubation in a galactose-based medium containing alginate beads without mat (Fig. 1B, 1C and supplementary text). The next most abundant ASV in G2.24 represents a sulfate-reducing bacterium (Desulfo-ASV1, 9%) of the order *Desulfovibrionales* (Fig. 1C). Low abundance organisms (>1% of ASVs) included *Chloroflexota, Thermoplasmatota*, and *Planctomycetota* (Data S1). Although some organisms were lost at different stages of cultivation, Desulfo-ASV1 and Loki-ASV2 were consistently observed together in varying ratios, supporting previous reports suggesting that Asgard archaea commonly form symbiotic relationships (*4, 5*).

To accurately classify Loki-ASV2 and Desulfo-ASV1 and investigate their capabilities and potential interactions, we used short- and long-read DNA sequencing of enrichment samples to assemble their complete genomes. The Loki-ASV2 circular genome contains 5,262,386 base pairs and encodes 4,544 predicted proteins, two sets of 16S/23S rRNA genes, and 46 tRNAs (table S1). Phylogenomic analyses based on 47 conserved marker genes with 112 reference genomes placed Loki-ASV2 as a sister lineage to the *Promethearchaeaceae* clade that includes *Promethearchaeum syntrophicum* MK-D1, *Candidatus* Harpocratesius repetitus FW102, and *Ca*. Lokiarchaeum ossiferum B35 (Fig. 1D, Data S3A-C). Synteny and pangenome analyses reveal an even distribution of the syntenic blocks, as well as orthologs of the four cultured *Promethearchaeaceae* (Fig. 1E, Data S2). The presence of apparent clusters of syntenic blocks in Loki-ASV2 may indicate that a few large genomic rearrangement events have occurred (Fig. 1E). For Loki-ASV2, we propose the name *Nerearchaeum marumarumayae* gen. nov., sp. nov., registered under the SeqCode (seqco.de/r:cjxbyjma) (*13*). The 4.1 Mb genome of Desulfo-ASV1 shows it is a novel bacterial genus and species within the family *Desulfovibrionaceae*, for which we propose the name *Stromatodesulfovibrio nilemahensis* gen. nov., sp. nov., registered under the SeqCode (seqco.de/r:cjxbyjma) (fig. S2, table S2, Data S2, Data S4A-C and supplementary text).

### *N. marumarumayae* cells show distinctive morphologies with a complex cell surface

Electron cryo-tomography (cryo-ET) was used to reveal the morphology and internal organization of cells in the enrichment cultures, focusing on the G2.24 high-enrichment culture (89% Loki-ASV2). Most cells had an overall coccoid morphology (avg. diameter 861 ± 109 nm), and many of these had substantial extensions that appeared either as chains of envelope vesicles (avg. diameter 100 ± 31 nm) (Fig. 2A-B, movie S1), or wide cytoplasmic tubules without distinct constrictions (Fig. 2A-F, movie S1). Some cells also appeared as multiple medium-sized cell bodies (200-500 nm in diameter) connected by long cytoplasmic tubules. Strings of interconnected cell bodies were observed, ranging up to 13 µm in length, with as many as five medium-sized cell bodies arranged in a row (e.g., fig. S1).

**Figure 2.**
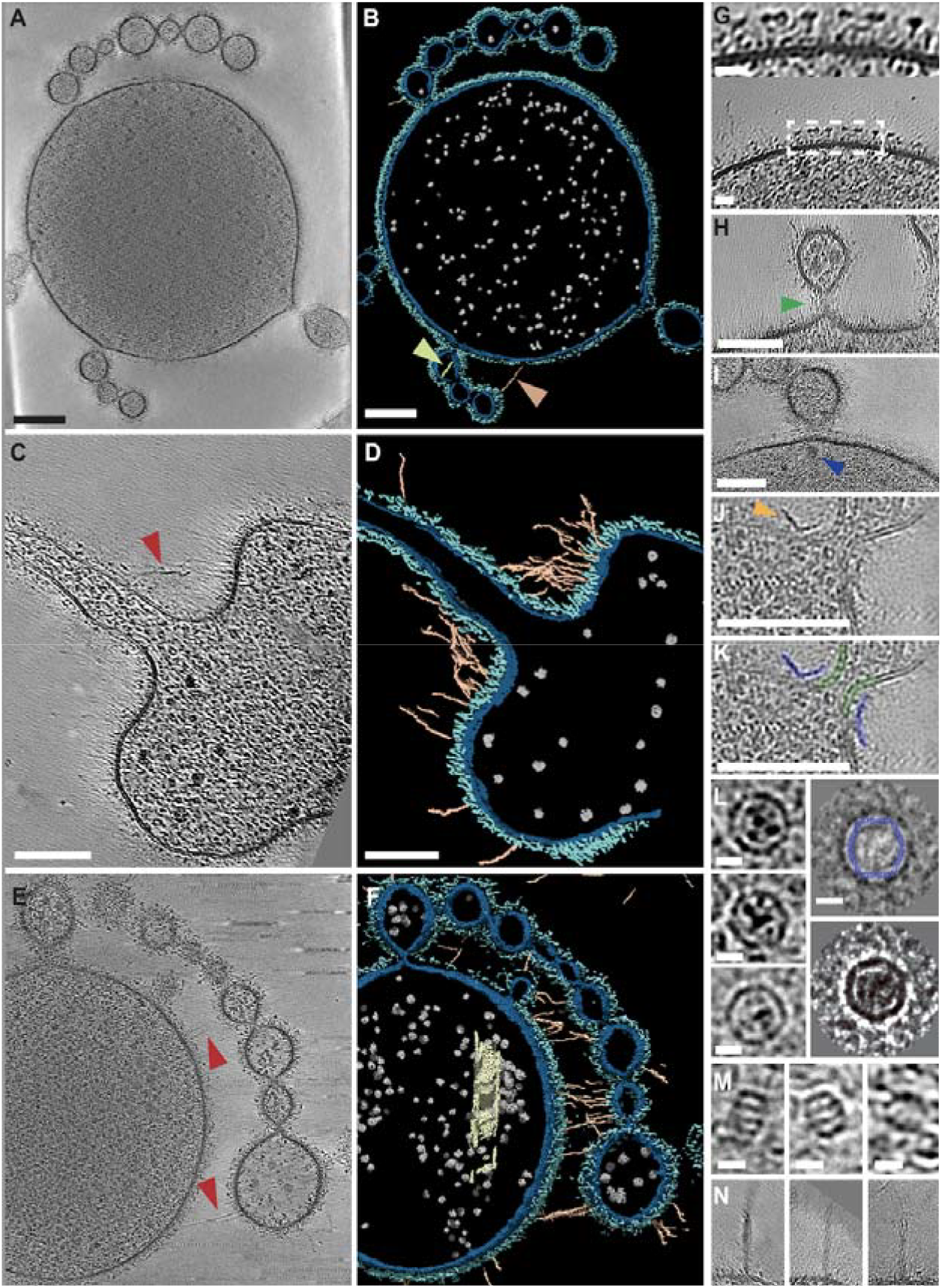
Structural features of *Nerearchaeum marumarumayae* cells. Electron cryo-tomography of *N. marumarumayae* cells. **(A)** Representative 2D slice through a 3D reconstructed tomogram, showing a cell body with smaller budded envelope vesicles that often appear as chains. **(B)** 3D Segmentation analysis of the tomogram in **A**, showing the identified inner membrane (blue), outer layer (teal), putative ribosomes (grey), extracellular fibers (light orange), and intracellular filament (light green). **(C)** Tomographic slice of a cell body with tubular extension and a large number of surface fibers. **(D)** 3D Segmentation analysis of the tomogram in C. **(E)** Tomographic slice of a cell body showing extensive budded vesicles that are connected to the main cell body through a network of extracellular fibers. **(F**) 3D Segmentation analysis of the tomogram in E, showing envelope vesicle detail and extracellular fibers connecting them to the cell body (indicated by red arrowheads where present in the selected 2D slice). The cell body also contains a large helical tube-like cytoplasmic structure (colored pale yellow). **(G)** Tomographic slice showing *N. marumarumayae* envelope detail; upper panel shows magnified view of the region in the dashed-box. **(H)** 2D tomographic slice showing a change in membrane density at the neck of a budding envelope vesicle (green arrow). **(I)** Example of a large diffused cytoplasmic density found at the neck base of budding envelope vesicles (blue arrow). **(J)** Detail of a continuous-membrane neck of an envelope vesicle, showing specific ‘L-shaped’ extracellular density (amber arrowhead), which in **(K)** is highlighted blue, and cell membrane in green **(L)**. Three example encapsulin-like cytoplasmic particles (left) and 2D class average of 13 such particles (lower-right panel). The upper-right panel shows the same 2D class average with a slice section of the *T. maritima* encapsulin cage structure (PDB: 7MU1) overlayed in blue. **(M)** Three example thermosome-like particles found in the cytoplasm. **(N)** Three example tower-like extracellular cell surface appendages. Scale bars 100 nm for **A**-**F** and **H-K** and 10 nm in **G** and **L-N**.

The coccoid cells and interconnected cell bodies revealed a highly distinctive cell envelope, consisting of a bilayer membrane decorated with a non-crystalline irregular outer surface (Fig. 2B, D, F, cyan). The outer surface possessed globular densities that extend 12 ± 1.3 nm from the cytoplasmic membrane, spaced by a lower-density region, appearing as irregular T-shaped structures (Fig. 2G). These striking morphologies and surface appearance contrast with the highly ordered protein S-layer array observed in most other archaea but are consistent with the other two *Promethearchaeaceae, Ca*. L. ossiferum and *P. syntrophicum* (*4, 5*). The G2.24 culture showed ∼70% of cells with these distinctive characteristics under cryoET (84 of 121 cells identified in 8 search maps). We deduce that these cells represent forms of *N. marumarumayae* Loki-ASV2.

Closer inspection of *N. marumarumayae* cell extensions showed that the cell body and chains of envelope vesicles (e.g., Fig. 2A-B, E-F) were connected via an indistinct surface association (e.g., Fig. 2H, I, green arrow) or a continuous membrane neck (e.g., Fig. 2J). These might represent different stages of envelope vesicle budding or distinct functions. A large globular, proteinaceous density (25 ± 1 nm), at a frequency of 18.8 %, was observed in the cell body, at the base of the tethered envelope vesicles (Fig. 2I, blue arrow). In some tomograms, at the neck-like connections, the membrane was flanked by a distinctive extracellular curved or L-shaped density (Fig. 2J-K). We also observed that the envelope vesicles (Fig. 2A, E) and tubular cytoplasmic extensions (Fig. 2C) were almost always attached to the main cell body through an additional network of extracellular fibers, as revealed by 3D segmentation analysis (Fig 2B, D, F, amber fibers). These fibers were up to several hundred nanometers long but are thinner (3 nm diameter) than archaella, without any obvious repeat, and there was no discernable density observed at the base, suggesting they are not assembled by a dedicated large molecular machine.

Within the main cell bodies, we did not observe the cytoplasmic actin-like filaments seen in *Ca*. L. ossiferum (*5*), despite the apparent complexity of *N. marumarumayae* cell extensions that presumably require complex organization of cytoskeletal structures. However, thin (4.5 nm wide) filaments were observed inside the interconnected cell bodies, especially near the junction of the cell body and interconnecting tubule (Fig. 2B). Interestingly, other novel subcellular structures were also observed including a wide helical tube-like structure of unknown identity (50 x 282 nm, Fig. 2F), encapsulin-like nanocages (Fig. 2L), thermosome-like complexes (Fig. 2M, 14 × 11 nm) and tower-like extensions (Fig. 2N, 9.5 ± 0.9 nm wide and 112 ± 33 nm long, membrane to tip). We identified 15 of the putative encapsulin nanocages and generated a low-resolution 2D average (Fig. 2L, right), which suggests these are mostly icosahedral (20 nm across) assemblies. These nanocages also showed internal electron-dense structures and the overall assembly matched well the structure of the mycobacterial (*14*) and *Thermotoga maritima* encapsulin nanocage (PDB: 7MU1) (Fig. 2L, top-right), which function in metal storage (*15*).

### *N. marumarumayae* proteins related to cell surface and internal structures

Structural and functional analysis of all predicted proteins encoded in the complete *N. marumarumayae* genome was performed to gain insights into the formation of distinctive subcellular and surface structures observed above. Initial gene functional assignment was based on their sequences and identified protein domains. We then predicted cellular location and potential for membrane association, and their 3D folded structures. A Foldseek database was then built for querying with existing protein structures (Fig. 3) (see materials and methods). With this approach, we initially identified proteins implicated in various cell biological and structural activities (Fig. 3A), revealing a mix of homologs conserved in bacteria, archaea and eukaryotes, including many of the ‘eukaryotic signature proteins’ (ESPs) known to be present in Asgard archaea (*3, 12,16*). For example, we identified sequence and structural homologs of eukaryotic endomembrane trafficking proteins such as ESCRTs, a Sec23/24-like protein (supplementary text), and small GTPases and their regulators containing longin and Rb/LC7 domains (Fig. 3A). These might be involved in forming the envelope vesicles observed (Fig. 2) and reflect ancestral mechanisms incorporated during the evolution of eukaryotic intracellular vesicle trafficking.

**Figure 3.**
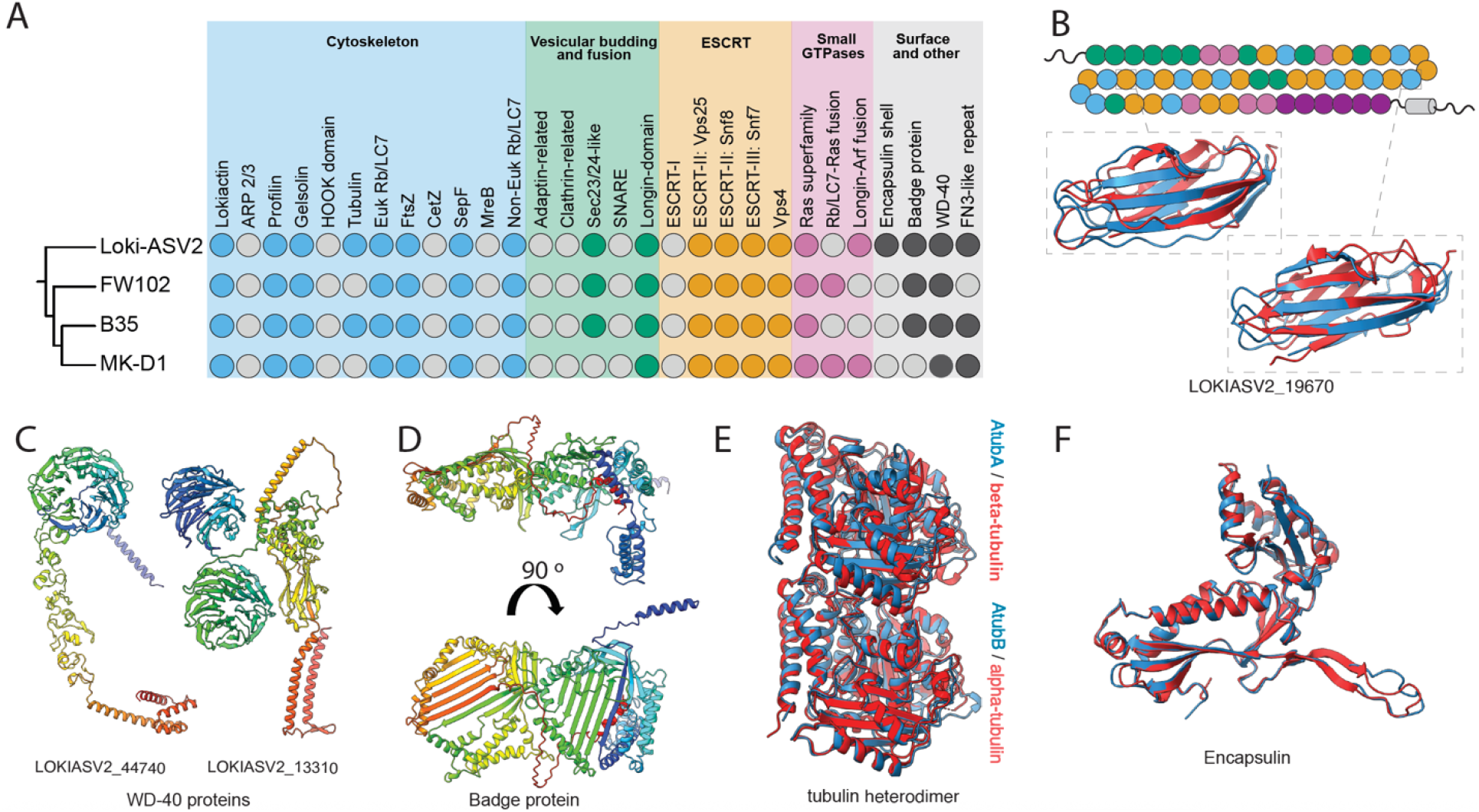
Conservation of selected homologs of eukaryotic and bacterial cell morphology and structural proteins in *Nerearchaeum marumarumayae*. **(A)** Schematic tree of *Promethearchaeaceae* cMAGs (left) wit corresponding table showing the presence (colored circles) or absence (grey circles) of the indicated proteins (right). **(B)** Schematic showing the organization of the 54 domains (circles) of LOKIASV2_19760 colored by five different fibronectin type-III (FN3) domain subtypes (Fig. S3). Structural alignment of LOKIASV2_19760 domains (blue) t experimental structures (red), showing similarity of domain 35 (middle) to a FN3 repeat from CbhA in *Clostridium thermocellum* (PDB: 3PE9, RMSD = 1.278 Å), and domain 20 (bottom) to an S-layer protein of *Deinococcus radiodurans* (PDB: 8CKA) (RMSD = 0.917 Å). **(C)** AlphaFold3 models of WD40 domain proteins with one WD40 domain (left) and two WD40 domains (right). **(D)** AlphaFold3 model of LOKIASV2_13140, with a ‘badge’ fold colored in a spectrum of red to violet by sequence/chain (N-to C-term.). AlphaFold3 models (blue) of **(E)** a hetero dimer of LOKIASV2_32170 (AtubB) and LOKIASV2_32180 (AtubA) aligned with the mammalian alpha/beta-tubulin heterodimer (PDB: 3J6E) (RMSD = 1.195 Å), and **(F)** LOKIASV2_08240 (blue) aligned with the *Thermatoga. maritima* encapsulin subunit (PDB: 7MU1) (RMSD = 0.843 Å).

Of the *N. marumarumayae* proteins 25.4 % were predicted to be extracellular membrane-bound proteins. Many of these were assigned as adhesin-like proteins, containing one or more fibronectin-like domains, a membrane-spanning domain, and predicted N-terminal signal sequences. For example, the largest *N. marumarumayae* protein (LOKIASV2_19760) comprises 5416 amino acids, showing some similarities to mammalian titin including 54 fibronectin-like domains (Fig. 3B, fig. S3), which could extend up to ∼250 nm from the membrane surface. We also searched for proteins with large extracellular globular domains that could be tethered to the cell surface seen by cryo-ET (e.g., Fig. 2G). Thirty-six WD40 beta-propeller proteins (ranging from 5-9 blades) were identified with many containing signal sequences and integral membrane domains. A number of these WD40 proteins also had fibronectin/adhesin domains between the membrane domain and the WD40 domain (e.g., Fig. 3C). WD40 domains occur in a wide-range of protein families (including COPII, and in sugar hydrolases like neuraminidase) and are typically associated with protein interaction scaffolds (*17*). WD40 domain proteins have been reported in archaeal and bacterial symbionts from coral reef microbiomes (*18, 19*). Other predicted surface-exposed proteins included a very large protein with a highly unusual and novel predicted fold that we referred to as a ‘badge’ protein due to its expected surface presentation and very large β-sheet domains (Fig. 3D). We were able to identify homologous proteins in all reported *Promethearchaeaceae* genomes (Fig. 3A). It is likely that such proteins contribute to the characteristic complex and irregular surface appearance of these organisms (Fig. 2G).

Although an extensive cytoskeleton was not obvious in the tomograms, we observed thin filament-like structures in the cytoplasm and budding vesicles (e.g., Fig. 2B, movie S1) and another unidentified wide tube-like structure (Fig. 2F). *N. marumarumayae* encodes cytoskeletal protein homologs, including an actin and its regulators profilin and gelsolin, and multiple tubulins (Fig. 3A). Two of the tubulins encoded by adjacent genes were predicted to form a heterodimer, which aligned closely to a mammalian alpha/beta-tubulin heterodimer from the microtubule lattice (Fig. 3E). Despite this, the *N. marumarumayae* tubulin sequences do not group specifically with eukaryotic alpha- or beta-tubulins in phylogenetic trees, and instead they affiliate with rare bacterial tubulin pairs (BtubA/B) found in *Prosthecobacter* spp (Fig. S5) that form mini-microtubules (*20*). Interestingly, one of the *N. marumarumayae* tubulins (LOKIASV2_32180) has a GTPase-inactive T7 loop, which is a characteristic of beta-tubulin for maintaining the GTP-bound heterodimer as the building block of eukaryotic microtubules (*21*). Furthermore, the *N. marumarumayae* tubulin pair is ∼84% identical to the recently characterized Asgard AtubA/B pair in *Ca*. L. ossiferum, which also forms mini-microtubules with as yet unknown function (*22*). *N. marumarumayae* is therefore likely to have cytoskeletal structures related to those seen in other Asgard archaea (*5, 22*), but their potential contributions to structures observed in *N. marumarumayae* cells remains to be seen.

*N. marumarumayae* encodes an encapsulin (Fig. 3F) and an encapsulin-associated ferritin-like protein (FLP), consistent with the appearance of distinct nanocages with internal electron dense structures observed in tomograms (Fig. 2L). This is the first time encapsulins have been observed in the *Promethearchaeaceae*. FLPs store iron and protect organisms against oxidative stress by converting ferrous to ferric iron (*15, 23*). Measurements of the pore water in Shark Bay mats have revealed that there are significant levels of free Fe^2+^ (*24*), suggesting a plausible role in the protection of *N. marumarumayae* against reactive oxygen species (ROS) generated by the Fenton reaction. *N. marumarumayae* also encodes a range of other enzymes that would protect cells against ROS (discussed below). This is likely important in microbial mats, which have dynamic diel fluctuation of oxygen levels due to oxygenic photosynthesis and aerobic respiration from surface cyanobacteria (*24-26*), as well as those at the subsurface utilizing infra-red light near the anaerobic lower layers (*27-29*). Other notable potential adaptations to the mat environment included a heliorhodopsin (possible UV tolerance - see supplementary text), and the chemotaxis and archaellum motility machineries.

### *N. marumarumayae* interacts with proximal bacteria

Several models of eukaryogenesis are based on inferred metabolic syntrophy between archaea and bacteria; however, direct contact between Asgard archaea and a syntrophic bacterium has not been observed. Our low magnification cryoEM images showed *N. marumarumayae* cells are often proximal or intertwined with diderm bacteria (Fig. 4A-B). High-resolution tomograms of the consortium revealed *N. marumarumayae* interacting with a diderm bacterium that has an LPS-based outer membrane (OM), a thin cell wall and inner membrane (IM) and polar flagellum, highly consistent with a cell envelope arrangement of a Gram-negative bacterium, expected to be *S. nilemahensis* (Desulfo-ASV1, 9% abundance in G2.24, Fig. 1C). Closer inspection revealed a series of thin tubular structures that originate in the bacterium, traverse the intercellular space and interact with the *N. marumarumayae* cell and its protrusions (Fig. 4C-D, arrowheads; movie S2). A flagellum and flagellar motor at the bacterial cell pole were clear (Fig. 4D, purple: flagellum; red: motor complex). Automated segmentation clearly distinguished between the thinner extracellular fibers that connect *N. marumarumayae* cell bodies and envelope vesicles (Fig. 2D, light orange filaments), and the intercellular nanotubes that extend from the bacterium to *N. marumarumayae* (Fig. 4D, pink filaments), confirming that they are two different appendages. Further inspection of the bacterial cell envelope revealed that there is an envelope-spanning protein complex at the base of each nanotube. This suggests that the archaea-bacteria interconnections are mediated by specific intercellular nanotubes.

**Figure 4.**
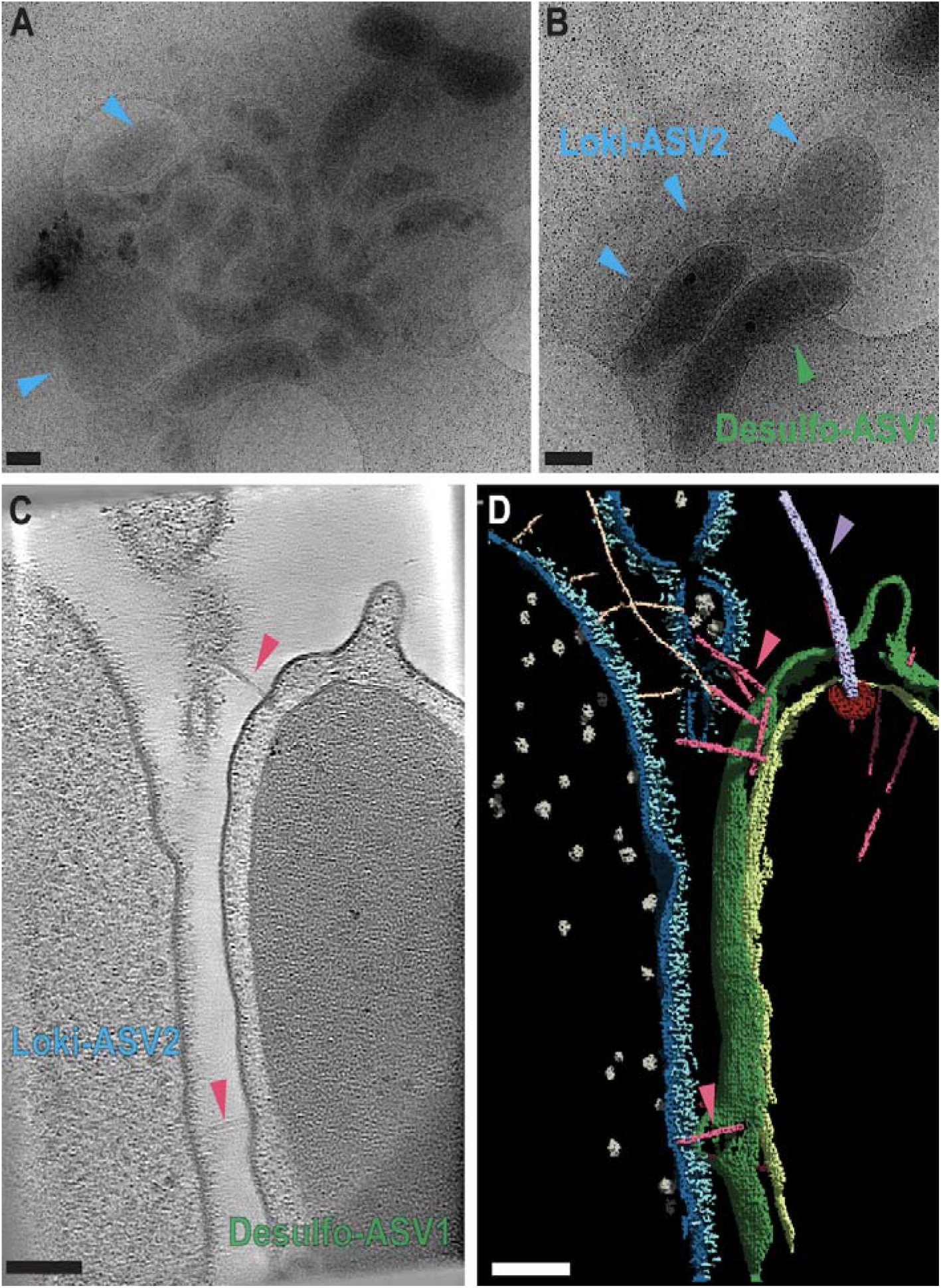
Direct interaction between *Nerearchaeum marumarumayae* and bacterial cells. **(A)** Low magnification cryo-TEM image of *N. marumarumayae* (Loki-ASV2) enriched cultures vitrified on an electro microscopy grid with carbon support. **(B)** Zoomed in cryo-TEM image of *N. marumarumayae* interacting wit bacteria (Desulfo-ASV1) cells via nanotubes (pink arrows). **(C)** 2D-slice through a 3D reconstructed tomogram of the interface between the cells shown in **B. (D)** Segmentation analysis of the tomogram shown in **C**, features are colored accordingly: *N. marumarumayae* cytoplasmic membrane (blue), outer layer (teal), putative ribosomes (grey), extracellular fibers (light orange), bacteria inner membrane (light green), bacterial outer membrane (green), bacterial sheathed flagella (purple), bacterial flagellar motor (red), and bacterial extracellular nanotube connecting *N. marumarumayae* cells (pink). Scale bars for panels **A, B** and **C, D** represent 500 nm and 100 nm respectively.

### Metabolic and syntrophic capacities of *N. marumarumayae* and *S. nilemahensis*

We undertook detailed genomic analyses to identify physiological and metabolic capacities of the two putative syntrophic partners in the context of the enrichment cultures and their microbial mat niche. For its nutrient assimilation and energy conservation strategies, *N. marumarumayae* is inferred to be a fermentative heterotroph (Fig. 5A, Data S5). It encodes the capacity to import and oxidize simple sugars, amino acids, and fatty acids; the product, acetyl-CoA, can be cleaved by acetyl-CoA synthetase to yield acetate and ATP. Another pathway for the generation of ATP by substrate-level phosphorylation in *N. marumarumayae* is through a partial, tetrahydrofolate-dependent methyl branch of the Wood-Ljungdahl pathway (WLP) for the catabolism of glycine, serine, and histidine, as recently suggested to be present in th common Asgard archaeal ancestor (*30*). The resulting reduced cofactors (NADH, reduced ferredoxin) can be used for biosynthesis, or re-oxidized using [FeFe]-hydrogenases (Fig. 5A, blue symbols, group A3 and B) that function as electron sinks with the release of H_2_ (*31*). The released H_2_ can be used in multiple ways. *S. nilemahensis* encodes a periplasmic uptak hydrogenase (group 1b [NiFe], Data S5) that couples the oxidation of H_2_ to sulfate and fumarat respiration (Fig. 5B). H_2_ uptake by *S. nilemahensis* would also help ensure that the H_2_ does not build up to inhibitory levels for *N. marumarumayae* (*4*). In the context of microbial mats, methanogenic archaea appear to dominate as consumers of H_2_ (and CO_2_) (*11, 26*). Thus, H_2_ appears to be an important energy currency in the mats involving multiple, diverse species.

**Figure 5.**
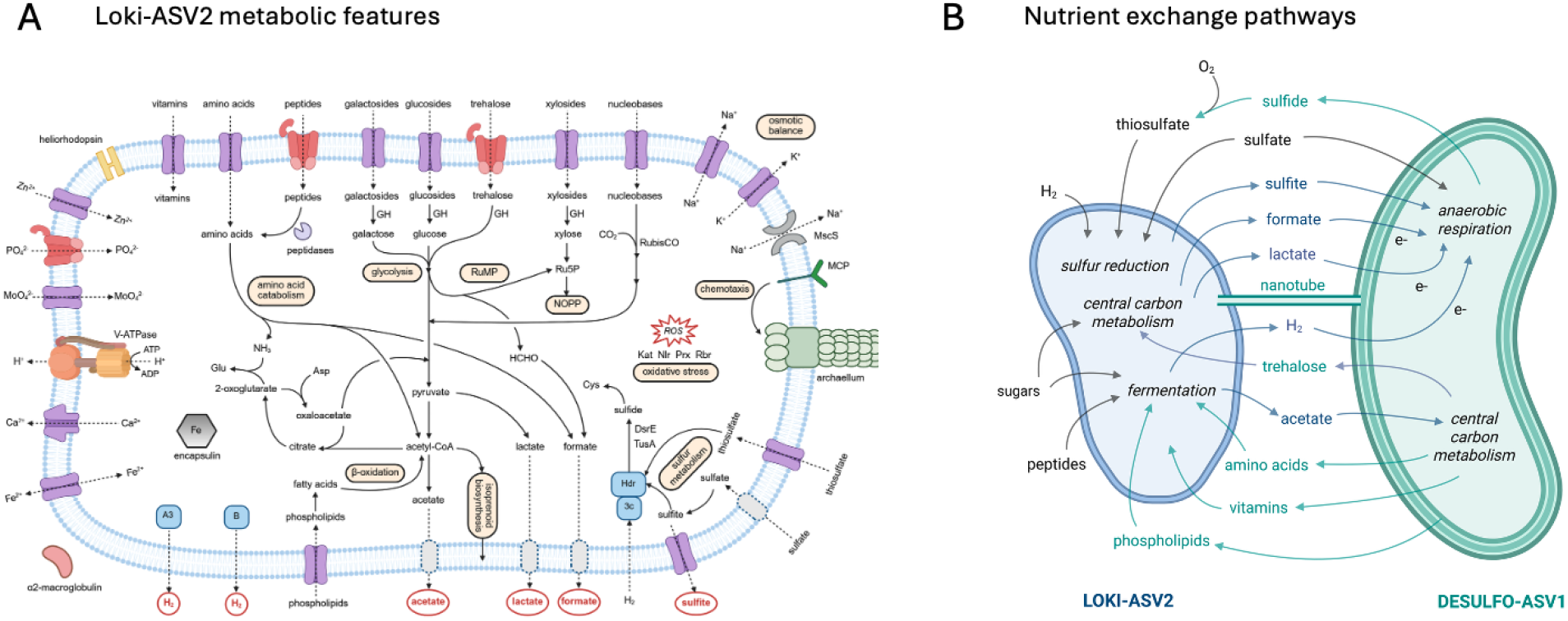
Metabolic networks in *Nerearchaeum marumarumayae* enrichments derived from microbial mats. **(A)** Genome-based metabolic reconstruction of the primary energy conservation pathways and selected nutrient and ion transporters in *N. marumarumayae*. H_2_, a key product of fermentation, can be oxidized by the group 3c [NiFe] hydrogenase, which is expected to be coupled to the reduction of sulfite and thiosulfate to sulfide (via TusA- and DsrE-bound persulfides) by cytosolic heterodisulfide reductase (Hdr). Sulfite is predicted to be generated b assimilatory sulfate reduction. Surplus sulfite can be exported by TauE transporters; thiosulfate is imported by TsuA transporters. *N. marumarumayae* metabolites predicted to be utilized by *S. nilemahensis* are circled in red; see mai text for further discussion. Abbreviations: GH, glycoside hydrolase; Kat, catalase; MCP, methyl-acceptin chemotaxis protein; MscS, mechanosensitive channel; Nlr, neelaredoxin; NOPP, non-oxidative pentose phosphate pathway; Prx, peroxiredoxin; Rbr, rubrerythrin; ROS, reactive oxygen species; RuMP, ribulose monophosphate pathway; V-ATPase, vacuolar/archaeal-type ATP synthase. **(B)** Proposed syntrophic interactions between *N. marumarumayae* and *S. nilemahensis*. Under this model, *N. marumarumayae* takes up trehalose, phospholipids, amino acids, vitamins, and thiosulfate from an extracellular pool of nutrients, and in turn releases metabolic by- products of potential benefit to *S. nilemahensis*, including H_2_, acetate, formate, lactate, and sulfite. Thiosulfate would be generated abiotically by chemical reaction of sulfide with oxygen in the microbial mat but would not be expected to be present in anoxic cultures. *S. nilemahensis* can synthesize the 20 essential amino acids, whereas the pathways for arginine, proline, phenylalanine, and tryptophan were not identified in *N. marumarumayae. S. nilemahensis* also has biosynthetic pathways for thiamine, cobalamin, pyridoxine, and riboflavin that were absent from *N. marumarumayae*. Figures created with BioRender.

*N. marumarumayae* may also utilize some of the H_2_, as it encodes a group 3c [NiFe]-hydrogenase-heterodisulfide reductase complex (*31*). We hypothesize that H_2_ oxidation is coupled to the reductive assimilation of sulfur (Fig. 5A), as *N. marumarumayae* lacks an identifiable sulfite reductase. Excess sulfite released by *N. marumarumayae* could also be utilized by *S. nilemahensis* for respiration (Fig. 5B) and would be especially beneficial to the latter as it obviates the need for sulfate activation initiated by expenditure of ATP (*32*).

In addition to the redox syntrophic exchanges described above, certain organic products of *N. marumarumayae* could be utilized by *S. nilemahensis* as energy and/or carbon sources (Fig. 5B), namely acetate and lactate generated fermentatively, and formate generated by a partial methyl branch of the WLP, as well as oxidation of formaldehyde, a by-product of the ribulose monophosphate pathway (*33*). In return, *S. nilemahensis* may be an important source of trehalose, fatty acids (from bacterial phospholipids), and certain amino acids and vitamins required but not synthesized by *N. marumarumayae* (Fig. 5B).

Finally, *N. marumarumayae* encodes two vacuolar/archaeal-type ATPase (V/A-ATPase). In the absence of identifiable ferredoxin-or NADH-dependent ion-translocation mechanisms, these V/A-ATPases may serve to hydrolyze ATP and drive proton export to generate a chemiosmotic gradient (Fig. 5A). Such a gradient may be used by secondary transporters for uptake of nutrients (sugars, amino acids and phospholipids) or sodium ion efflux via antiporters (Fig. 5A) that may help deal with the fluctuating salinity experienced in the microbial mats (*24*) (*34*). *N. marumarumayae* also encodes a primary transporter for trehalose uptake (Fig. 5A), which could serve as a compatible solute against high salinity in the mats, or as a carbon source. Release of trehalose, such as via cell lysis in mats (including from sulfate-reducing bacteria or cyanobacteria; (*35, 36*), may be utilized by *N. marumarumayae* (Fig. 5B). We speculate that the uptake of compatible solutes might have evolved in mat systems on an early Earth where oceans were hypersaline (*37*).

## Conclusions

The syntrophic and physical interactions described here between an Asgard archaeon and a sulfate-reducing bacterium from a modern microbial mat environment aligns with hydrogen-sulfur syntrophy models (*7, 38*) for the origin of eukaryotes in ancient microbial mats. Our findings suggest that H_2_ and formate produced by an Asgard archaeon are used for energy by a sulfate-reducing bacterium; acetate is used for anabolic functions in the bacterium; and sulfite produced by the Asgard archaeon is transferred to the bacterium, where it serves as a substrate for energy conservation by sulfite respiration. Conversely, organic compounds generated by the bacterium may be utilized by the archaeon, supporting its growth and fermentative metabolism.

The budded envelope vesicles, extracellular fibers, and substantial intracellular tube-like structures of *N. marumarumayae* cells further support structural models of eukaryogenesis where cellular extensions of Asgard archaeal ancestors led to more intimate and interdependent microbial associations to promote syntrophy, eventually leading to engulfment and the evolution of specialized cell structures and organelles (*4, 39*). In the dynamic, structured environments of microbial mats, continually fluctuating stresses may have driven ancient symbioses like the one we describe here, to enhance survival, providing a platform for eukaryogenesis. Microbial mats may indeed be the ‘microbial village that helped raise a eukaryote’ (*40*).

## Etymology

Ne.re.ar.chae′um. Gr. masc. n. *Nereus* (Nηρεύς) in Greek mythology, ancient sea god (described as the ‘Old Man of the Sea’) and father of the Nereides, female spirits of sea waters, in reference to both the coastal marine origin of this microorganism, and the antiquity of microbial mats and stromatolites; N.L. neut. n. *archaeum*, an archaeon; N.L. neut. n. *Nerearchaeum*, an archaeon named for Nereus; *marumarumayae*. ma.ru.ma.ru.ma’yae. derived from the Indigenous language of the Malgana people of Shark Bay where the microbial mats and stromatolites are sourced, meaning ‘ancient home’, a reference to stromatolites being of ancient origin in Earth’s geological and biological history; *marumaru*, many nights, old, ancient; *maya* camp, home. N.L. gen. n. *marumarumayae*, of the ancient home.

*Stromatodesulfovibrio* stro.ma.to.de.sul.fo.vi′bri.o. Gr. fem. gen. n. *stromatos*, mat, layer, in reference to the layers found in microbial mats. N.L. masc. n. *Desulfovibrio*, a bacterial genus name. N.L. masc. n. *Stromatodesulfovibrio*, a *Desulfovibrio*-like bacterium from a microbial mat. *nilemahensis*. ni.le.mah.en’sis. after Nilemah, place in Western Australia where the microbial mat is located; N.L. masc. adj. -*ensis. nilemahensis*, from Nilemah.

## Acknowledgments

We are grateful to Malgana language expert Kymberley Oakley, Elder Auntie Pat Oakley, Malgana Aboriginal Corporation, and Malgana Elders for helping find a suitable name for Loki-ASV2 and for granting permission to use the words *marumaru maya* from the language of the Malgana people of Gathaagudu (Shark Bay). We also acknowledge support from Malgana rangers and coordinators in the field. All fieldwork and sampling were conducted using valid permits issued by the Department of Biodiversity, Conservation and Attractions (DBCA), Western Australia. We acknowledge the support of the MWAC Structural Biology Facility (https://doi.org/10.26190/4KQF-M552) and UNSW Computing Cluster Katana (https://doi.org/10.26190/669x-a286) for access to computational resources. CryoET data were collected at the Ian Holmes Imaging Center (Bio21, University of Melbourne).

## Author contributions

S-JN established the high abundance enrichments, genomic and phylogenetic analyses, protein sequence- and structure-based identification, and their functional analyses. MDJ acquired, analyzed, interpreted, and visualized cryoET data that led to detailed cell biology descriptions including the proposed physical cell-cell interactions observed. TJW did the metabolic reconstructions, pathway and syntrophy analyses, and interpretation. JM established and optimized the high abundance enrichments, produced initial MAGs that facilitated taxonomic placement, and performed genome annotations, phylogenetic, and metabolic analyses. XV-C reconstructed the cMAGs and did the detailed genome analyses, annotation, phylogenies, interpretation, and description of new genera/species. FIM established the enrichment cultures and conducted the initial metagenomic analyses through taxonomic classification, metabolic reconstructions, and analysis of encoded cell biology machinery of the enriched organisms. KR did the computational protein structure prediction, searches, and curation. MP did genomic DNA analyses and Nanopore sequencing. BP contributed to cryoET analyses and segmentation. DCS contributed to cryoET analyses and segmentation. KAM planned, oversaw and interpreted the computational protein structure analyses and enrichment. IGD oversaw metabolic and cellular analyses and contributed to interpreting overall results. DG planned and oversaw the structural biology project component, performed and interpreted cryoET. BPB initiated and oversaw the overall project and contributed to interpreting the results. S-JN, MDJ, TJW, JM, XV-C, MP, KR, KAM, IGD, BPB, DG wrote the manuscript with input and edits from all authors.

## Data and materials availability

The *N. marumarumayae* and *S. nilemahensis* MAGs and raw metagenomic data are available under BioProject PRJEB83094. Raw amplicon sequencing data of the enrichment cultures are available under BioProject PRJEB87679. The in silico structural proteome data for Loki-ASV2 are available at https://github.com/keiran-rowell-unsw/Loki-ASV2_in_silico. Other data are available in the main text or the supplementary materials.

