## Supplemental text and figures for "An Asgard archaeon from a modern analog of ancient microbial mats"

Nobs et al., 2025.

**The PDF file includes:**

Materials and Methods

Supplementary Text

Figs. S1 to S7

Tables S1 to S3

References (*1-139*)

**Other Supplementary Materials for this manuscript include the following:**

Movies S1 to S2

Data S1 to S8

**Contents**

**Materials and Methods**………………………………………………………………...…….…..3

**Supplementary Text**………………...…………………………………………………………..11

**Supplementary Figures**

**fig S1.** Low-magnification cryoET 2-dimensional (2D) projection images of Loki-ASV2 enrichment cultures derived from microbial mats. ………………………….…………..17

**fig. S2**. *Desulfovibrionaceae* concatenated protein tree. ……………….…………….....18

**fig. S3**. Structural phylogeny of repeated fibronectin type-III-like domains in LOKIASV2_19760. ………………………………………………………………………..……….….….......19

**fig. S4.** Phylogeny of eukaryotic-like tubulins rooted to artubulins …………….……...20

**fig. S5.** AlphaFold3 model of LOKIASV2_32180 (AtubA) and LOKIASV2_32170 (AtubB)................…………………………………………………………………..…....21

**fig. S6.** AlphaFold3 model of LokiASV2_08240……………………………………….22

**fig. S7**. Phylogeny of heliorhodopsin showing placement of Loki-ASV2 _25410…..…23

**Supplementary Tables**

**table S1.** Genome assembly details for *Nerearchaeum marumarumayae* Loki-ASV2 and the reference circularized *Promethearchaeaceae* MAGs. ………………………………24

**table S2.** Genome assembly details for *Stromatodesulfovibrio nilemahensis* Desulfo-ASV1…………………………………………………………………………………..…25

**table S3.** Loki-ASV2 AtubAB alignment to mammalian alpha/beta tubulin……………26

**Supplementary Movies and Data descriptions.** …………………………..…………..………27

**Materials and Methods**

**Microbial mat sampling.** Cultures were established from samples of smooth microbial mats (Fig. 1) at the Nilemah tidal flats in Shark Bay (*Gathaagudu*), Australia (26˚27’336’’S, 114˚05’762’’E) collected on 15^th^ March 2019. These samples, designated SB19, were collected as previously described (*10, 25, 26*). Conditions at the time of sampling were temperature 34 °C, salinity 70 PSU, solar radiation 22.5 MJ/m^2^. Samples were collected during low tide at just below water level and all samples were collected and handled with sterile instruments. Samples were stored at 4 °C during transport back to the laboratory. DNA extraction from mat samples, 16S rRNA gene amplicon and metagenomic sequencing of environmental DNA, was described previously (*10*).

**Initial establishment of cultures from microbial mats.** Sections of microbial mat (2 g) were taken between ca. 12–18 mm below the mat surface (covering the anoxygenic zone, (*25*) were dispersed and in 120 mL of anaerobic medium in serum flasks that were sealed with butyl stoppers. A range of carbon sources (xylan, glucose, cellulose, yeast extract) at 0.05% w/v were employed as an initial enrichment strategy, and relative abundance assessed with 16S rRNA gene amplicon sequencing (described below) after three months incubation. Initial cultures were established in media with cellulose, yeast extract or glucose (0.05% w/v) supplemented with casamino acids (0.05% w/v) in a basal salt medium reflective of Hamelin Pool seawater (50.7 g/L NaCl, 13.3 g/L MgSO_4_·7H_2_O, 7.23 g/L MgCl_2_·6H_2_O, 2.7 g/L CaCl_2_·2H_2_O, 1.4 g/L KCl) (*41*). Media were also supplemented with SL10 trace elements solutions (1 mL/L) (*42*), vitamin 10 solution (3 mL/L) (*43*), and antibiotics; ampicillin (100 µg/mL), kanamycin (50 µg/mL) and streptomycin (100 µg/mL). Subsequent transfers following initial inoculation contained the same medium but were supplemented with 4 g/L of homogenized microbial mat, sterilized by autoclaving. All cultures were grown under anaerobic conditions generated by gas sparging (80:20% v/v N_2_:CO_2_). Media was further reduced with Na_2_S·9H_2_O (0.3 g/L) and L-cysteine (0.3 g/L) and pH was adjusted to 7.5. Anaerobic conditions were monitored with resazurin (100 µg/mL) and cultures were incubated at 30 °C without shaking. DNA extractions on initial cultures were performed with the DNeasy PowerBiofilm Kit (Qiagen).

**Establishment and monitoring of high abundance Asgard enrichments.** Starting from a cellulose-based medium, further enrichment strategies were tested and optimized over successive generations (Data S6). Galactose (0.05% w/v) was used as a primary carbon source due to the presence of multiple galactosidases encoded in Asgard archaeal MAGs. In addition to supplementing media with autoclaved microbial mat, culturing attempts were made either without microbial mat samples or using alginate beads replacing autoclaved microbial mat (see below for details). Monitoring consisted of analysis of Asgard archaea relative abundance via 16S rRNA gene amplicon sequencing of cultures (described below) two months after establishment, and cultures with Asgard archaeal enrichment of ca. 20% or higher were transferred every three months. Earlier culture generations were also monitored with light microscopy on fixed cells and samples containing high biomass were assessed for levels of Asgard archaea relative abundance via 16S rRNA gene amplicon sequencing. High enrichment was achieved following 6 months of incubation in media as follows: galactose (0.05% w/v), casamino acids (0.05 % w/v), Hamelin Pool basal salt solution (50.7 g/L NaCl, 13.3 g/L MgSO_4_·7H_2_O, 7.23 g/L MgCl_2_·6H_2_O, 2.7 g/L CaCl_2_·2H_2_O, 1.4 g/L KCl), SL10 trace elements (1 mL/L), Na_2_SeO_3_ (4 mg/L), Na_2_WO_4_·2H_2_O (4 mg/L), ampicillin (100 µg/mL), kanamycin (50 µg/mL), streptomycin (100 µg/mL), tetracycline (100 µg/mL) and alginate beads (prepared as described below). When included, tryptophan was added at a concentration of 0.0005% w/v). Sparging and addition of cysteine and sodium sulfide (as described above) were done after the addition of the beads.

Alginate beads were prepared using 2% (w/v) sodium alginate solution and 0.2 M CaCl_2_. To reduce the presence of oxygen, the alginate and CaCl_2_ solution were moved into an anaerobic chamber (gas mix) while hot, following autoclaving, and allowed to cool to room temperature in the chamber. Beads were prepared by dropping small aliquots of the alginate solution into the CaCl_2_ solution with a 25G needle and syringe. For the beads to form, the CaCl_2_ solution was stirred gently by hand at the same time. The CaCl_2_ solution containing alginate beads was then sealed in an anaerobic jar and incubated at 150 rpm at room temperature for a further 30 min. The alginate beads were drained using an autoclaved tea strainer under anoxic conditions and ten beads were transferred per serum bottle.

**DNA extraction and 16S rRNA gene amplicon sequencing.** DNA was extracted from cultures using the NEB Monarch Genomic DNA Purification Kit following manufacturer’s instructions for purification from Gram-positive Bacteria and Archaea with the following modifications. Cells were harvested through centrifugation for 10 min at 20,000 × *g*. After cell lysis and RNase treatment, cellular debris and sediment were separated from cell lysate through centrifugation at 20,000 × *g* for 10 min. The supernatant was transferred to new tubes and then gDNA binding buffer was added. DNA concentration was quantified with a Qubit 4.0 fluorimeter using the dsDNA HS kit and quality was assessed with a Nanodrop spectrophotometer UV absorption spectrum. Amplicon sequencing was performed on the extracted DNA at the Ramaciotti Centre for Genomics on an Illumina Mi**S**eq v3 2×300 bp sequencing run using universal primer set 926F (5′AAACTYAAAKGAATTGRCGG3′) and 1392wR (5′ACGGGCGGTGWGTRC3 ′) that cover all three domains of Archaea, Bacteria and Eukarya (*44*). After quality control including low-quality read removal, data was annotated for taxonomic relative abundance using the DADA2 pipeline (*45*) against the SILVA database 138.1.

**Long-read sequencing and base calling.** DNA from Asgard archaea enrichment cultures was extracted as described above. DNA was quantified using the Qubit 1× dsDNA HS kit and a Qubit Flex Fluorometer (Invitrogen). DNA fragment length was determined via a Genomic DNA ScreenTape (200–60,000 bp range) and TapeStation 4150 platform (Agilent). DNA extracts from three enrichment cultures were pooled (SB19_G3_3b_4_24, SB19.G2.18_1_24, GluSJM_2_11_23) and purified using AMPure XP beads (1:1 ratio) (Beckman Coulter). The library was prepared using the Ligation Sequencing Kit V14 (SQK-LSK114, Oxford Nanopore Technologies, ONT) with an input of 504 ng, and the following modifications. The end prep thermocycler was set for longer time to improve the recovery due to low DNA input (20 °C for 5 min, 65 °C for 5 min to 20 °C for 7 min, 65 °C for 7 min), and all bead purification elution steps, including the initial bead clean-up, were performed at 37 °C instead of room temperature to improve the retention of HMW DNA. The final library was washed with long fragment buffer (LFB). Flow cells (R10.4.1, FLO-MIN114) were primed with Bovine Serum Albumin (BSA) (50 mg/mL) (Invitrogen) and loaded with 201.6 ng (~27 fmol) of library. Sequencing was performed on the ONT GridION platform for 72 hours. Reads were base-called offline with Dorado v0.7.3 (https://github.com/nanoporetech/dorado) and the corresponding super accuracy model v5.0.0 in duplex mode (dna_r10.4.1_e8.2_400bps_sup@v5.0.0). Sequencing generated 22.95 Gbp (~95% pass), 8.35 M reads and N50 of 4.09 kbp.

**Long read assembly**. Long-read data were assembled with metaFlye v2.9.4-b1799 (*46*) and provided with the --nano-corr option. To improve Loki-ASV2 assembly quality, contiguous contigs in Flye’s assembly graph belonging to Loki-ASV2 were manually extracted in Bandage v0.8.1 (*47*) and the ONT reads mapped back to them with minimap2 v2.28-r1209 (*48*) with lr:hq mode. Mapped reads were extracted with SAMtools v1.13 (*49*). Reads were then filtered with SeqKit v2.5.1 (*50*) to remove reads shorter than 500 bp and with a Q<15 before reassembly with metaFlye. This resulted in a single circular chromosome. The Loki-ASV2 chromosome was then polished with short reads obtained from shotgun metagenomes. First, Polypolish-default v0.6.0 (*51*) was used and the output was further processed with Pypolca-careful v0.3.1 (*52, 53*). Reorientation of Loki-ASV2 and the reference genomes on the COG1474 (CDC6-related/ORC1-type DNA replication protein) was initially attempted with Dnaapler v0.8.0 (*54*) in archaea mode but it was unable to detect the full-length of the COG1474 resulting in the beginning of the chromosomes being set within the gene. Therefore, known COG1474 sequences from the reference genomes were used with Dnaapler in custom mode.

**Genome annotation**. Genome annotation was performed with DFAST v1.3.1-c1dc63e (*55*) with all the options for pseudogene prediction deactivated due to overestimations in poorly described Asgard lineages. Prodigal v2.6.3 (*56*) was used as the gene-caller, CRT v1.2 (*57*) for CRISPR detection, and tRNAscan-SE v2.0.12 (*58*) for the detection of tRNAs. In addition to the default DFAST database, functional annotation included blastn and blastp searches against CARD v3.2.9 (*59*) and VFDB 20240716 (*60*) databases; HMM-based searches against TIGRFA M v15.0 (*61*), Pfam v37.0 (*62*), and dbCAN3 HMM profiles v12 (*63*), and RPS-BLAST against COG (2020 release) (*64*), KOG (*65*) and CDD 20240330 (*66*). Pseudogene detection was deactivated in DFAST due to the inflated pseudogene detection rate in our dataset. In addition to the default Barrnap 0.9 (<https://github.com/tseemann/barrnap>) and tRNAscan-SE, overall ncRNAs were predicted with infernal v1.1.4 (*67*) with Rfam 14.10 (*68*) (options --cut_ga --rfam --nohmmonly -Z <2×genome size>) due to the limited detection of rRNA genes by Barrnap in the *Promethearchaeaceae* circular genomes.

Additional functional annotation of protein coding genes were performed with eggNOG-mapper v2.1.2 (*69*) against the default eggNOG v5.0.2 (*70*) and the “novel_fam” databases (*71*). InterProScan v5.68-100.0 (*72*) was run with options --disable-precalc --goterms --iprlookup --pathways, and all the supported databases and tools: AntiFam v7.0 (*73*), CDD v3.20 (*74*), Coils v2.2.1 (*75*), FunFam v4.3.0 (*76*), Gene3D v4.3.0 (*77*), HAMAP 2023_05 (*78*), MobiDB-lite v2.0 (*79*), NCBIfam v4.0 (*80*), PANTHER v18.0 (*81*), Pfam v37.0 (*62*), Phobius v1.01 (*82*), PIRSF v3.10 and PIRSR v2023_05 (*83*), PRINTS v42.0 (*84*) ProSitePatterns v2023_05 and ProSiteProfiles v2023_05 (*85*), SFLD v4 (*86*), SignalP v4.1 (*87*), SMART v9.0 (*88*), SUPERFAMILY v1.75 (*89*), and TMHMM v2.0c (*90*). KEGG-based annotations were carried out in anvi’o v8-dev (*91*) with adaptive threshold adjustment (*92*) and the --include-stray-KOs option. Hydrogenase sequences were classified using the HydDB online webserver. Default parameters were used, unless otherwise stated.

**Phylogenomic analyses.** Initial taxonomic assignment of the circularized MAGs was performed with GTDB-Tk v2.4.0 (*93*) against the GTDB r220 (*94*), and the output used to define the taxa included in the phylogenomic analyses. For the Loki-ASV2 tree, analysis was circumscribed to o__Syginarchaeales, with o__Helarchaeales as outgroup. Representative genomes from both orders were filtered based on the number of detected marker genes (at least 50%, 112 total reference genomes). The marker genes from the standard 53 archaeal markers were also filtered out to remove all those rarely present in the selected genomes resulting in 47 markers being retained for phylogenetic analysis. Marker sequences from the reference genomes and Loki-ASV2 were extracted with GTDB-Tk’s align command. Sequences from each marker were aligned with MAFFT L-INS-i v7.526 (*95*) and trimmed with ClipKit v2.3.0 (*96*) with a gap threshold of 50% (--gaps 0.5). Three methods were used for the phylogenomic analysis: concatenated proteins, partitioned analysis and multispecies coalescent (MSC) model supertree. (i) Concatenated protein: trimmed alignments were concatenated, and trees were built with IQ-TREE v2.3.3 (*97*) with the LG+F+R8 substitution model (-m MFP) (*98*). Support values were calculated with 1000 ultrafast bootstrap replicates (UFBoot2) with nearest neighbor interchange optimization (--bnni) (*99*). (ii) Partitioned tree: the concatenated tree was used as input for IQ-TREE with partitions and substitution models defined based on the markers. Support values were calculated with 1000 ultrafast bootstrap replicates (UFBoot2) with nearest neighbor interchange optimization (--bnni) (*99*). UFBoot2 replicates were performed by resampling partitions and then resampling sites within resampled partitions (--sampling GENESITE) (*100*). (iii) MSC tree: built with ASTER v1.16 with the ASTRAL-hybrid algorithm using ML trees from all individual markers as input (*101*). Trees for each individual marker were built based on the trimmed alignments with IQ-TREE -B 1000 --bnni with the same substitution models as their corresponding partitions in (ii).

The phylogenomic analyses of Desulfo-ASV1 were performed in a similar manner but with the following modifications. The tree was limited to f__Desulfovibrionaceae with f__Desulfonauticaceae as an outgroup. Marker genes were selected from the standard 120 SCG markers from GTDB, resulting in a final set of 115 SCGs and 336 reference genomes. The substitution model for the concatenated protein tree was LG+F+I+R10.

**Average amino acid identity (AAI).** Average amino acid identity (AAI) of Loki-ASV2 was calculated with CompareM v0.1.2 (<https://github.com/dparks1134/CompareM>) against the reference cMAGs. Similarly, AAI between Desulfo-ASV1 and the closest genomes was based on the phylogenomic analyses.

**Comparative genomics of cMAGs/CIRCOS plot.** Orthologous proteins across the four *Promethearchaeaceae* cMAGs were predicted with Broccoli v1.3 (*102*). Syntenic blocks were detected with SynChro (January 2015 release) (*103*), as mapping-based methods struggle with more distant genomes, e.g. across different genera. GC skew was calculated with SkewIT (commit: 4f096fc) (*104*). GC content across the genomes was calculated every 20 bp with a sliding window of 100 bp with SeqKit v2.5.1 (*105*). Default parameters were used for all tools unless otherwise stated.

**Focused search for cytoskeletal and morphological proteins.** The predicted protein sequences for all four closed *Promethearchaeota* genomes were annotated using the Galaxy Australia instance of InterProScan (v. 5.59-91.0) with Pfam, TIGRFAM, PANTHER, SUPERFAMILY, Gene3D, Hamap, SMART, PRINTS, PrositePatterns, FunFam, PIRSF, PrositeProfiles selected. Proteins that matched PFAM or InterPro entries of proteins of interest were pulled out and underwent further assessment for annotation. Further assessment consisted of: a protein homology search with DIAMOND (*106*) (v2.1.7) in sensitive mode against the NCBI nr database (221v5) (*107*) filtered for eukaryotic sequences (taxid: 2759) and another iteration against the entire database excluding unclassified archaea (taxid:93506) and *Promethearchaeati* (taxid: 1935183), an orthology search using the Galaxy Australia instance of eggNOG Mapper (v. 5.0.2) with DIAMOND seed orthologues, and FoldSeek (*108*) of ESMFold -predicted *in silico* structures against the Swiss-Prot, Proteome and PDB100 databases using software versions detailed below. Queries with hits to unrelated proteins were removed as well as those with bitscores below 100. Queries with FoldSeek bitscores between 100-200 were further investigated and removed based on poor structural homology. The taxonomy of the seed-orthologue in EggNOG Mapper as well as the taxonomy of DIAMOND and FoldSeek hits were used to determine if the protein was eukaryotic- or prokaryotic-like. Low probability matches across multiple searches with poor structural matches were removed from the analyses. The list of annotated cytoskeletal and morphological proteins is located in Data S7.

**Mapping metabolic pathways based on annotated genomes.** Metabolic traits were identified based on the annotated genomes of *N. marumarumayae* and *S. nilemahensis* combined with manual assessment of the veracity of gene functional assignments (*109-111*). To determine the presence or absence of an encoded enzyme catalyzing a specified reaction, the predicted protein sequence(s) carrying out the reaction from the genome annotation were individually submitted to ExPASy BLAST (using the “UniProtKB/Swiss-Prot only” option) (*112*). To verify its functional assignment, proteins were required to show ≥ 35% sequence identity to an experimentally verified protein with the unambiguous metabolic function in the ExPASy BLAST database. If this identity threshold was not reached, InterProScan (*113*) results for all proteins were searched to identify functional domains and potential for membrane or intra- or extracellular locations, as the basis for assigning a catalytic function if there were no known ambiguities. In several cases of remaining ambiguity, proteins were described as hypothesized or proposed to catalyze a particular reaction considering gene synteny and neighboring steps in the pathway where possible. Glycoside hydrolase (GH) families were classified according to CAZy (Carbohydrate-Active enZymes) (*114*). Protein sequences that were identified as hydrogenases based on catalytic domains were classified further using the hydrogenase classifier HydDB (*115*).

**Computational 3D protein structure predictions.** Protein structures for all 4544 protein coding sequences in the closed Loki-ASV2 genome were predicted using several different contemporary AI protein structure prediction codes, depending on the constraints of the protein sequence. ESMFold (*116*) v1.0.3 (git commit ID [2b36991](https://github.com/facebookresearch/esm/commit/2b369911bb5b4b0dda914521b9475cad1656b2ac)) was run in batch mode across a .FASTA file containing all protein coding amino acid sequences from the Loki-ASV2 MAG detailed above. ESMFold predicted 4372/4544 protein structures, until the sequence lengths (>880 amino acids) produced out-of-memory errors as the model inference demands exceeded available GPU RAM (40 GB NVIDIA A100). The remaining protein sequences were predicted with AlphaFold (*117*) v2.3.2 (commit ID [f251de6](https://github.com/google-deepmind/alphafold/commit/f251de6613cb478207c732bf9627b1e853c99c2f)), with the exception of three proteins longer than 3,000 amino acids (index #s 975, 1976, 3246) and two proteins that had consistent execution errors along the AlphaFold 2.3 pipeline (index #s 3250, 3449). AlphaFold2 was run with access to the standard databases from the download_all_data.sh script, which were stored on local NVMe SSDs of the server for quick execution of standard bioinformatics search and alignment algorithms. The five remaining proteins after ESMFold and AlphaFold2 execution were predicted using AlphaFold3 (*118*) at DeepMind’s [alphafoldserver.com](http://alphafoldserver.com/). Protein #1976 was the largest sequence in the Loki-ASV2 proteome at 5416 amino acids. Due to the Alphafold server’s 5,000 amino acid limit, the structure was predicted in two portions and then reassembled manually in PyMol (*119*).

The ‘top’ result from the above protein folding codes were used to assemble an *in silico* proteome structure set, this is either the one result from ESMFold, or the ‘rank 0’ prediction from AlphaFold 2 and 3. A Python script, modified from a PyMol wiki example (*120*) was used to render an image of each protein within this *in silico* proteome in standard orientation and colored by AlphaFold-style pLDDT residue prediction confidence metrics. All raw prediction output and scripts for processing are available at the GitHub repository associated with this project (<https://github.com/keiran-rowell-unsw/Loki-ASV2_in_silico>). The final set of predicted 3D protein structures are uploaded to ModelArchive (<https://modelarchive.org/doi/10.5452/ma-6k8yb>) which provides hosting and a stable DOI for theoretical biomolecular structure models.

**Structure based homology searches.** Foldseek (*108*) (tag [9-427df8a](https://github.com/steineggerlab/foldseek/tree/9-427df8a), commit ID 427df8a) was used to find related homologues to the *in silico* proteome based not on sequence identity but on structural identity, using the 3Di alphabet which encodes nearest-residue positionally information using a variational autoencoder. The 20 symbols representing the local structural factors at each residue are then aligned in much the same way as traditional bioinformatics alignment methods. In this case foldseek easy-search was used with default settings (with the exception of repeats in the titin homologue, where the --alt-ali was used to match domains with the predicted structure). Foldseek databases were the standard set (UniProt, UniProt50, UniProt50-min, Proteome, Swiss-Prot, ESMAtlas30, PDB100, CATH50, ProstT5, Mgnify-2022-05) downloaded via the foldseek databases command. Additionally, the more recently released BFMD, and BFVD databases were downloaded from the mirror ([foldseek.steineggerlab.workers.dev/](https://foldseek.steineggerlab.workers.dev/)) set up in the Steinegger lab at Seoul National University, Republic of Korea.

For annotation purposes, a script was written to create a table that contains the top foldseek match of each Loki-ASV2 *in silico* protein from either PDB100 or Swiss-Prot due to their focus on higher quality annotations. Additionally, an ‘all vs all’ set of foldseek matches were created, through a script that ran foldseek easy-search for each protein against all the aforementioned foldseek databases and collated them into a ‘_all_foldseek_dbs.tsv’ file for each protein. Lastly, human-friendly foldseek .HTML reports were created with the ‘–format-mode 3’ flag. The ‘all vs all’ reports took slighty over a month to generate using a single 64-thread AMD EPY 7313 16-Core CPU, processing against foldseek databases on HDD network attached storage across a 1GbE link. All foldseek output files are also available at the project repository (<https://github.com/keiran-rowell-unsw/Loki-ASV2_in_silico>).

**Reciprocal FoldSeek search.** The *in silico* proteome was made into a customizable and searchable database using the foldseek createdb option to find putative structural homologs of experimentally validated structures. A curated set of pdb files (Data S8) were used as queries against the custom database and searched using foldseek easy-search with .HTML reports generated. Queries included but are not limited to; vault protein, dynamin, spectrin, and crescentin. Matches with bitscores over 100 were checked for their corresponding annotations from the initial genome annotation, focused protein search and structure-based homology searches. New matches are found in Data S8.

**Protein structural analyses.** Protein structures featured in this paper were remodeled using the AlphaFold 3 webserver with default settings, except for AtubA (LOKIASV2_32180) and AtubB (LOKIASV2_32170), which were folded together with a monomer of each. Alignments were made on ChimeraX (v1.8) (*121*) using MatchMaker with default settings (bb chain pairing using the Needleman-Wunsch alignment algorithm), unless otherwise stated (Table S3). The encapsulin shell protein was aligned to a monomer from *Thermotoga maritima* encapsulin (*15*); PDB:7MU1). Root mean square deviations (RMSD) of aligned pairs of proteins represent the pruned atom pairs. A new structure-based homology search was conducted on these new models as outlined above against the PDB, SwissProt, and Proteome databases. Domains of the model of LOKIASV2_19760 were outputted as individual PDB files and a structural alignment was made using the FoldMason online webserver (<https://search.foldseek.com/foldmason>) (*122*) validated structures of fibronectin-type 3 (FN3) domain and immunoglobulin-like domain (Ig) from human titin (PDB: 8OT5 and 1G1C, respectively). The amino-acid and 3Di alignments were concatenated and a partitioned maximum-likelihood phylogeny was made with IQ-TREE (v2.2.2.5) based on the methods and substitution matrix described previously (*123*).

### Electron cryotomography data collection. For freezing, samples were first mixed with 10 nm colloidal gold beads (Sigma-Aldrich, Australia), which were precoated with 1% BSA. The samples were then pipetted onto glow-discharged R2/2 Quantifoil holey carbon grids (Quantifoil Micro Tools GmbH, Jena, Germany), and excess liquid was removed by back blotting before plunging into liquid ethane. The entire freezing process was performed inside the Vitrobot chamber (FEI Thermo Fisher Scientific) under 100% humidity. Grids were imaged using a Titan Krios G4 cryo-EM, operating at 300 kV acceleration voltage, and equipped with a Gatan energy filter and a K3 Summit direct detector. Tilt series images were collected in movie mode using a dose-symmetric scheme at two different pixel sizes, 3.39 Å and 1.603 Å, using Tomography 5 software version 5.14 (Thermo Fisher Scientific) at a tilt range of −51° to 51° in 3° increments, with defocus values of -6 μm and -4 μm, respectively. Data were collected with a cumulative dose of ~120 e-/Å² per tilt series.

### Electron cryotomography data processing. Frames were initially motion-corrected using MotionCor3 (*124*) and then aligned with the IMOD software package (*125*) integrated within Scipion (*126*). The aligned tilt series were then reconstructed at either bin 8 or bin 4 using IMOD or Tomo3D (*127*). To enhance interpretability, tomograms were denoised using Cryo-CARE (*128*). For 3D segmentation, in some cases, fiducials were erased from the tilt series using Fidder (<https://github.com/teamtomo/fidder.git>) prior to tomogram reconstruction.

### Subtomogram analysis. Encapsulin-like particles were extracted as sub-tomograms from 8x-binned (12.82 Å/pix) CTF corrected tomograms using Dynamo (v1.1.546) (*129*). An initial model of the encapsulin-like particle was generated by averaging extracted sub-tomograms without alignment. For sub-tomogram averaging the extracted particles were aligned for one round with three iterations.

### Tomogram segmentation. To better visualize the tomograms, 3D volumes were segmented using Dragonfly software, v2022.2 (<https://www.theobjects.com/dragonfly/index.html>). Before segmentation, built-in filters were applied to improve clarity. This was followed by the manual segmentation of 20 to 30 slices from individual tomograms, which were then used as input for neural network training using the U-Net architecture with a 2.5D input dimension (7 slices) (*130*). The trained model was subsequently used to segment the entire tomogram, with manual corrections implemented where necessary. The color schemes used were as follows: OM (blue), non-crystalline surface layer (sky), filaments (cream), cytoplasmic thin tube (green) and ribosome (grey). Built-in functions were used to generate 2D images and 3D movies.

**Supplementary Text**

**Asgard archaea could be enriched from microbial mats in Shark Bay.** Hamelin Pool in Shark Bay, Western Australia, is characterized by three dominant types of microbial mats – smooth, pustular, and colloform (*131*). For the present study, smooth microbial mats sampled from the upper-intertidal region of the Nilemah region of Hamelin Pool were used as the starting inoculum as previous analyses of these mat types via shotgun metagenomic sequencing revealed the presence of diverse Asgard archaeal MAGs (*10, 11*). Preliminary 16S rRNA gene amplicon sequencing indicated that Asgard archaea were present at 3.7% relative abundance in these smooth mats (*132*). A number of analyses of the microbial composition of microbial mats in Shark Bay have been conducted (e.g. (*26*)), however the Asgard archaea were then classified as Marine Benthic Group B. Preliminary MAGs assembled from shotgun sequencing of enrichment cultures indicated that Glycoside Hydrolase Family 3 was a frequently occurring enzyme family encoded in draft lokiarchaeal genomes which led to the decision to trial xylan and cellulose-based media alongside the glucose-based media as a standard carbon source. Enrichment of the Asgard archaea consistently reached up to 20-35% after 3-6 months incubation over successive transfers, however, higher enrichments were not attained until further conditions were tested as outlined in the main text.

***N. marumarumayae* and *S. nilemahensis* represent new genera and species.** Long-read assemblies were able to recover circularized MAGs (cMAGs) of the main representatives in the cultures: Loki-ASV2 and Desulfo-ASV1. The Loki-ASV2 cMAG contains 5,262,386 bp, with a GC content of 34.0%, encoding 4,544 predicted proteins, two copies of each rRNA (16S, 23S and 5S), and 46 tRNAs (table S1). Concatenated and partitioned phylogenomic analyses place Loki-ASV2 within *Promethearchaeceae*, as sister to the clade formed by all the named *Promethearchaeceae* genera: *Promethearchaeum*, *Ca.* Lokiarchaeum, and *Ca.* Harpocratesius (Fig. 1, Data S2-S5). The supertree also places Loki-ASV2 with the previously indicated genera, but its position is uncertain (Data S5). AAI between Loki-ASV2 and the representative cMAGs indicate values well within the range of the same family but distinct genera (54.5–55.5%, Data S2). The 16S rDNA sequences of Loki-ASV2 are the most divergent of the *Promethearchaeaceae* cMAGs with 90.6–91.2% identity with the other representatives.

The assembled Desulfo-ASV1 cMAG is 4,134,807 bp long, with a 61.0% GC, encoding 3,779 proteins, two full rRNA operons (16S, 23S and 5S), and 51 tRNAs (table S2). Preliminary phylogenetic analysis with GTDB-Tk, place Desulfo-ASV1 within the family *Desulfovibrionaceae*. In all phylogenetic trees, Desulfo-ASV1 is part of a well-supported clade with *Desulfobaculum*, *Desulfocurvus*, *“Desulfovibrio” ferrophilus*, and the placeholder genus g__UBA6814 (Fig. S2, Data S6-S8). In both the concatenated and partitioned trees, Desulfo-ASV1 appears as sister of g__UBA6814, while the Desulfo-ASV1-g__UBA6814 clade is sister to the *Desulfobaculum*-*Desulfocurvus*-*“Desulfovibrio” ferrophilus* lineage. However, the position of Desulfo-ASV1 with respect to the other *Desulfovibrionaceae* is uncertain in the supertree. AAI values between Desulfo-ASV1 and all the clade representatives’ range within 56.9–58.9% (Data S2). *“Desulfovibrio cavernae”* H1MT (AJ621885.1) has the closest 16S rRNA gene sequence of any named organism (valid or not) with 96.7% identity. While the 16S rRNA gene sequence may indicate Desulfo-ASV1 belongs to the same genus as *“Desulfovibrio cavernae”*, this was considered not validly published and pre-date the *Desulfovibrio* sensu lato reclassification efforts in the genomic era (*133*).

Phylogenetic analyses, AAI calculations, and lack of 16S rRNA gene matches with any nomenclatural type indicate that both Loki-ASV2 and Desulfo-ASV1 constitute the first representatives of two new genera. Thus, under the SeqCode, we propose the names of *Nerearchaeum marumarumayae* Loki-ASV2 gen. nov. sp. nov. within the family *Promethearchaeaceae*, and *Stromatodesulfovibrio nilemahensis* Desulfo-ASV1 gen. nov. sp. nov. within the family *Desulfovibrionaceae*. The new genera and species have been registered in the SeqCode with the following registry ID: [seqco.de/r:cjxbyjma](https://registry.seqco.de/registers/r:cjxbyjma).

**Structural analyses of LOKIASV2_25730** **suggests it is Sec23/24-like.** While the initial accession-based search also indicated that Loki-ASV2 and B35 uniquely contained a sec23/24 trunk domain containing protein, cross-referencing with the pangenome orthologous clusters indicated it was also shared with FW102 (Fig. 1, OG_2173) which was only identified to have the sec23/24 zinc finger domain. The best homology match annotation indicates it could be more like a circularly permuted Ras-like GTPase (cpRAS) (*134*). Further structural assessment was undertaken to suggest it is Sec23/24-like. ESMFold structures of OG_2173 were aligned to each other and to cpRAS (AF-Q75J93) on PyMOL (v 3.0.3) which indicated OG_2173 did not contain the GTPase motif at the N-terminal of the protein. RMSD values of the protein from *Ca.* L. ossiferum B35 and *P. syntrophicum* MK-D1 to Loki-ASV2 were 2.504 and 2.037 respectively, while LOKIASV2_25730 aligned to cpRAS was 4.103. LOKIASV2_25730 was then folded with AlphaFold3 containing a zinc ion and compared to Sec23/24 (PDB:1M2V) using ChimeraX (see above). Alignment resulted in an RMSD of 7.799 Å to Sec23 (PDB: 1MV2, chain A) between 505 pairs with no iterations. Upon comparison, LOKIASV2_25730 was notably missing the gelsolin-like domain but contained a Cys4 zinc-finger despite having no annotation as such~~.~~ This missing gelsolin domain is consistent with a recent report of Asgard Arf GTPases (*135*).

**Loki-ASV2 tubulins are predicted to form an alpha/beta tubulin-like heterodimer**. To assess the phylogenetic relationship of the two *N. marumarumayae* tubulins with eukaryotic tubulins, the sequences were added to a previous collection of curated tubulin sequences (*3*). The phylogeny (fig. S4) suggests the LOKIASV2_32180 and LOKIASV2_32170 are related to eukaryotic tubulins, though they form a sister lineage to bacterial tubulin A (BtubA) and BtubB, respectively. These bacterial proteins were believed to have originated via horizontal gene transfer from an early eukaryote or Asgard archaeon to *Prosthecobacter* spp (*20, 22*). To be consistent with these studies, we named the *N. marumarumayae* proteins AtubA (LOKIASV2_32180) and AtubB (LOKIASV2_32170). The T7 loop sequences of these *N. marumarumayae* tubulins, which are normally involved in activating GTPase activity to promote microtubule disassembly, further supported eukaryotic-like stable hetero-dimerization; AtubB contains the catalytic glutamic acid residue (E251) conserved in alpha-tubulin and AtubA contains the conserved catalytically-inactive lysine residue (K248) as in beta-tubulin, which together result in a stable alpha/beta pair that forms the primary unit of polymerization and depolymerization in microtubules.

The *N. marumarumayae* AtubAB heterodimer structural model (AlphaFold3), colored by pLDDT in ChimeraX using a b-factor (confidence) color palette (alphafold), is shown along with iPTM and PTM values in fig. S4. Table S3 shows statistics of the 3D structures aligned to eukaryotic alpha/beta-tubulin (PDB 3J6E). Since MatchMaker alignment with default settings on ChimeraX aligns the best-fitting chains, the ‘ss’ chain pairing mode had to be used to obtain the RMSD when both pairs were included in the alignment (overall RMSD of 2.745 Å). However, alignment with default settings of the AtubA monomer with the alpha/beta-tubulin heterodimer indicated that chain B (beta-tubulin) was the best match for AtubA, with an RMSD of 1.950 Å across all 419 atom pairs. Finally, specifying monomer alignments between AtubB and 3J6E chain A (alpha-tubulin) provided an RMSD of 2.313 Å across 411 atom pairs (table S3).

**Encapsulin shell protein has strong structural matches to *T. maritima* encapsulin**. The encapsulin shell protein (LOKIASV2_08240) was remodeled (AlphaFold3 server) which produced a highly confident prediction of the protein (Fig. 3C, fig. S6). Structure-based homology search with FoldSeek against SwissProt and PDB databases indicates high similarity to the shell protein in the hyperthermophilic bacterium *Thermotoga*. The top hit in SwissProt was to type 1 encapsulin shell protein from *Thermotoga* sp. (strain RQ2) (accession:B1L7S2, bitscore: 1304); the top hit in the PDB database was to a Cryo-EM structure of an encapsulin form *T. maritima* (PDB: 7KQ5, bitscore: 1267). The default alignment with matchmaker on ChimeraX had an RMSD of 0.843 Å between 241 pruned atom pairs, and 1.383 Å between all 262 atom pairs to the encapsulin shell protein in *T. maritima* (PDB: 7MU1).

**A** **heliorhodopsin has a putative role in stress response**. A heliorhodopsin (LOKIASV2_25410) in Loki-ASV2 was identified via InterProScan (IPR041113) and clustering of the cMAGs indicated it is a shared feature with *Ca.* L. ossiferum B35 and *P. syntrophicum* MK-D1. We undertook a phylogenetic assessment to check if it might be a schizorhodopsin. The sequence was added to an existing alignment (*136*) using MAFFT (v7.481) and trimmed using ClipKit (v2.3.0) with kpic-based trimming. A maximum-likelihood phylogenetic tree was created with IQ-TREE (v2.2.2.5) using ModelFinderPlus (*137*) and 10000 ultrafast bootstrap replicates. The heliorhodopsin from *N. marumarumayae* clusters with other heliorhodopsins found in Asgard archaea (fig. S7) as opposed to the schizorhodopsins found in Asgard archaea (*136*). While its precise function is unknown, the Loki-ASV2 heliorhodopsin may be a sensor that upregulates DNA photolyase in UV damage repair (*138*) which would be necessary in the high UV environment of Shark Bay (*10*). Schizorhodopsins, which act as light-driven inward H^+^ pumps (*136, 139*), were not found in Loki-ASV2 even though they are encoded in other Asgard archaeal metagenomes from microbial mats (*11*).

**Taxonomy**

**Description of *Nerearchaeum* gen. nov.**

Ne.re.ar.chae′um. Gr. masc. n. *Nereus* (Νηρεύς) in Greek mythology, ancient sea god (described as the 'Old Man of the Sea') and father of the Nereides, female spirits of sea waters, in reference to both the coastal marine origin of this microorganism, and the antiquity of microbial mats and stromatolites; N.L. neut. n. *archaeum*, an archaeon; N.L. neut. n. *Nerearchaeum*, an archaeon named for Nereus.

Genus defined based on the phylogenomic analysis of 47 single copy marker genes, and average amino acid identity calculations against known members of the family *Promethearchaeaceae*. The type species is *Nerearchaeum marumarumayae*.

This taxon is registered in the SeqCode in the following register list: seqco.de/r:cjxbyjma.

**Description of *Nerearchaeum marumarumayae* sp. nov.**

ma.ru.ma.ru.ma’yae, derived from the indigenous language of the Malgana people of Shark Bay where the microbial mats and stromatolites are sourced, meaning ‘ancient home’, a reference to stromatolites being of ancient origin in Earth’s geological and biological history; *marumaru,* many nights, old, ancient; *maya* camp, home. N.L. gen. n. *marumarumayae,* of the ancient home.

The type material is the metagenome assembled genome Loki-ASV2 obtained from enrichment cultures of smooth microbial mats (stromatolites) at the Nilemah tidal flats in Shark Bay (*Gathaagudu*), Australia. The MAG consists of a single circularized contig of 5.26 Mbp, containing two copies of each 16S, 23S and 5S genes and 46 tRNAs (22 unique: 20 standard tRNA-aa plus tRNA-iMet and tRNA-SeC). The GC content of this MAG is 34.0%.

Based on functional genomic analysis, predicted to be a fermentative acetogenic heterotroph; predicted metabolic pathways, processes, and features as follows: Glycolysis (Embden-Meyerhof-Parnas pathway) that includes pyrophosphate-dependent 6-phosphofructokinase; gluconeogenesis; pentose phosphate pathway (non-oxidative only); ribulose monophosphate (RuMP) pathway; truncated tricarboxylic acid cycle (comprises pyruvate:ferredoxin oxidoreductase, ATP citrate synthase, aconitate hydratase, isocitrate dehydrogenase, 2-oxoacid:ferredoxin oxidoreductase, malate dehydrogenase). Hydrogenases include two [FeFe] hydrogenases (group A3); one [FeFe] hydrogenase (group B); one [NiFe] hydrogenase (group 3c), the latter inferred to form a complex with heterodisulfide reductase. Glycoside hydrolases, including β-galactosidases, β-glucosidases, and β-xylosidases. Peptidases, including aminopeptidases, carboxypeptidases, dipeptidases, and gingipain-like peptidases. Organic substrates include glucose, galactose, xylose, mannonate, galacturonate/d-glucuronate, glycerol, myo-inositol, alcohols, aldehydes, alanine, glutamate, asparagine, serine, glycine, histidine, lysine, leucine, isoleucine, valine, cysteine, fatty acids. Partial methyl branch (oxidative) of the Wood-Ljungdahl pathway (tetrahydrofolate-dependent), leading to formate. Ammonia assimilation by glutamine synthetase. ABC transporters for uptake of trehalose/maltose, nucleobases, peptides, phosphate, manganese, and thiamine. Other transporters for uptake of sugars, amino acids, phospholipids, nucleobases, 5'-deoxyadenosine, vitamins (including riboflavin, pantothenate), thiosulfate, phosphate, zinc, iron(II), cobalt, molybdate, potassium, chloride. Other transporters for efflux of sulfite, arsenate, arsenite, manganese, cadmium, cobalt, magnesium, zinc, copper, calcium, and sodium. Two vacuolar/archaeal-type (V/A-type) ATP synthases. ATP generation by substrate-level phosphorylation using acetyl-CoA synthetase. Synthesis of glutamate, glutamine, aspartate, asparagine, serine, glycine, alanine, threonine, cysteine, methionine, lysine, tyrosine, leucine, isoleucine, valine, histidine, putrescine, isoprenoid phospholipids. Sulfatases for hydrolysis of sulfate esters. Assimilatory sulfate reduction pathway, missing sulfite reductase (heterodisulfide reductase may perform this step). Sulfur relay proteins include TusA, DsrE family, and rhodanese-like protein. Defense against oxidative stress by catalase-peroxidase, neelaredoxin (superoxide dismutase), desulfoferredoxin (superoxide reductase), rubrerythrin, rubredoxin, peroxiredoxin. Defense against nitrosative stress by hydroxylamine reductase, nitric oxide reductase. Capsulin nanocompartment containing ferritin-like cargo protein (involved in oxidative stress protection). Motility and chemotaxis using archaella. Heliorhodopsin (possible sensor involved in UV damage repair). Alpha2-macroglobulin (possibly involved in intercellular interactions).

This taxon is registered in the SeqCode in the following register list: seqco.de/r:cjxbyjma

**Description of *Stromatodesulfovibrio* gen. nov.**

[Stro.ma](http://stro.ma/).to.de.sul.fo.vi′bri.o. Gr. fem. gen. n. *stromatos*, mat, layer, in reference to the layers found in microbial mats. N.L. masc. n. *Desulfovibrio*, a bacterial genus name. N.L. masc. n. *Stromatodesulfovibrio,* a *Desulfovibrio*-like bacterium from a microbial mat.

Genus defined based on the phylogenomic analysis of 115 single copy marker genes, and average amino acid identity calculations against known members of the family *Desulfovibrionaceae*.

This taxon is registered in the SeqCode in the following register list: seqco.de/r:cjxbyjma

**Description of *Stromatodesulfovibrio* *nilemahensis* sp. nov.**

ni.le.mah.en’sis, after Nilemah, place in Western Australia where the microbial mat is located; N.L. masc. adj. -*ensis. nilemahensis,* from Nilemah*.*

The type material is the metagenome assembled genome Desulfo-ASV1 obtained from enrichment cultures of smooth microbial mats (stromatolites) at the Nilemah tidal flats in Shark Bay (*Gathaagudu*), Australia. The MAG consists of a single circularized contig of 4.13 Mbp, containing two copies of each 16S, 23S and 5S genes and 51 tRNAs (22 unique: 20 standard tRNA-aa plus tRNA-fMet and tRNA-SeC). The GC content of this MAG is 61.0%.

Based on functional genomic analysis, predicted to be an anaerobic heterotroph; predicted metabolic pathways, processes, and features as follows: Glycolysis (Embden-Meyerhof-Parnas pathway); gluconeogenesis; pentose phosphate pathway (both oxidative and non-oxidative); incomplete tricarboxylic acid cycle (succinate dehydrogenase absent); synthesis of trehalose, glycogen, all 20 amino acids, glycine betaine, thiamine, cobalamin, riboflavin, pyridoxine. Hydrogen oxidation by periplasmic [NiFe] hydrogenase (group 1b). Anaerobic respiration using sulfate, sulfite, cysteate, isethionate, fumarate, dimethylsulfoxide. F-type ATP synthase; vacuolar (V-type) ATPase. Rnf-Mrp Na^+^/H^+^ antiporter complex. CO oxidation by carbon monoxide dehydrogenase. Organic substrates include glucose, acetate, formate, lactate, citrate, phosphoglycolate, glycerol, ethanolamine, phosphonoacetate, (*R*)-1-hydroxy-2-aminoethylphosphonate, 2-aminoethylphosphonate, peptides, glutamate, aspartate, asparagine, serine, leucine, isoleucine, valine, diaminopropionate, alcohols, aldehydes. Ammonia assimilation by glutamine synthetase. Multiple peptidases, including aminopeptidases, carboxypeptidase, dipeptidases. ABC transporters for uptake of peptides, glutathione, branched-chain amino acids, lysine/arginine/ornithine, glutamine, tyrosine, glycine betaine, purine, sulfonates, phosphonates, phosphate, molybdate, tungstate, zinc. Other transporters for uptake of sulfate/sulfite, thiosulfate, isethionate, acetate, lactate, citrate, monocarboxylates, C4-dicarboxylates, glucose, ammonium, glutamate, tyrosine, glycine betaine, proline, choline, ectoine, biotin, pantothenate, phosphonates, zinc, magnesium, cobalt, ferrous iron, potassium. Other transporters for efflux of amino acids, iron, nickel, cobalt, cadmium, zinc, copper, arsenite, calcium, sodium. Sodium-pumping rhodopsin. Nitrogen fixation by nitrogenase. Defense against oxidative stress by cytochrome *bd* ubiquinol oxidase, superoxide dismutase, catalase-peroxidase, neelaredoxin, thioredoxin reductase, rubredoxin-oxygen oxidoreductase, peroxiredoxin. Defense against nitrosative stress by hydroxylamine reductase. Bacterial microcompartment (for ethanolamine catabolism). Type IVc pilus (Tad) secretion system. Type II secretion system (T2SS). Two-partner secretion (TPS) system (FhaB/FhaC-like). Motility and chemotaxis using flagella.

This taxon is registered in the SeqCode in the following register list: seqco.de/r:cjxbyjma

**Supplementary Figures**

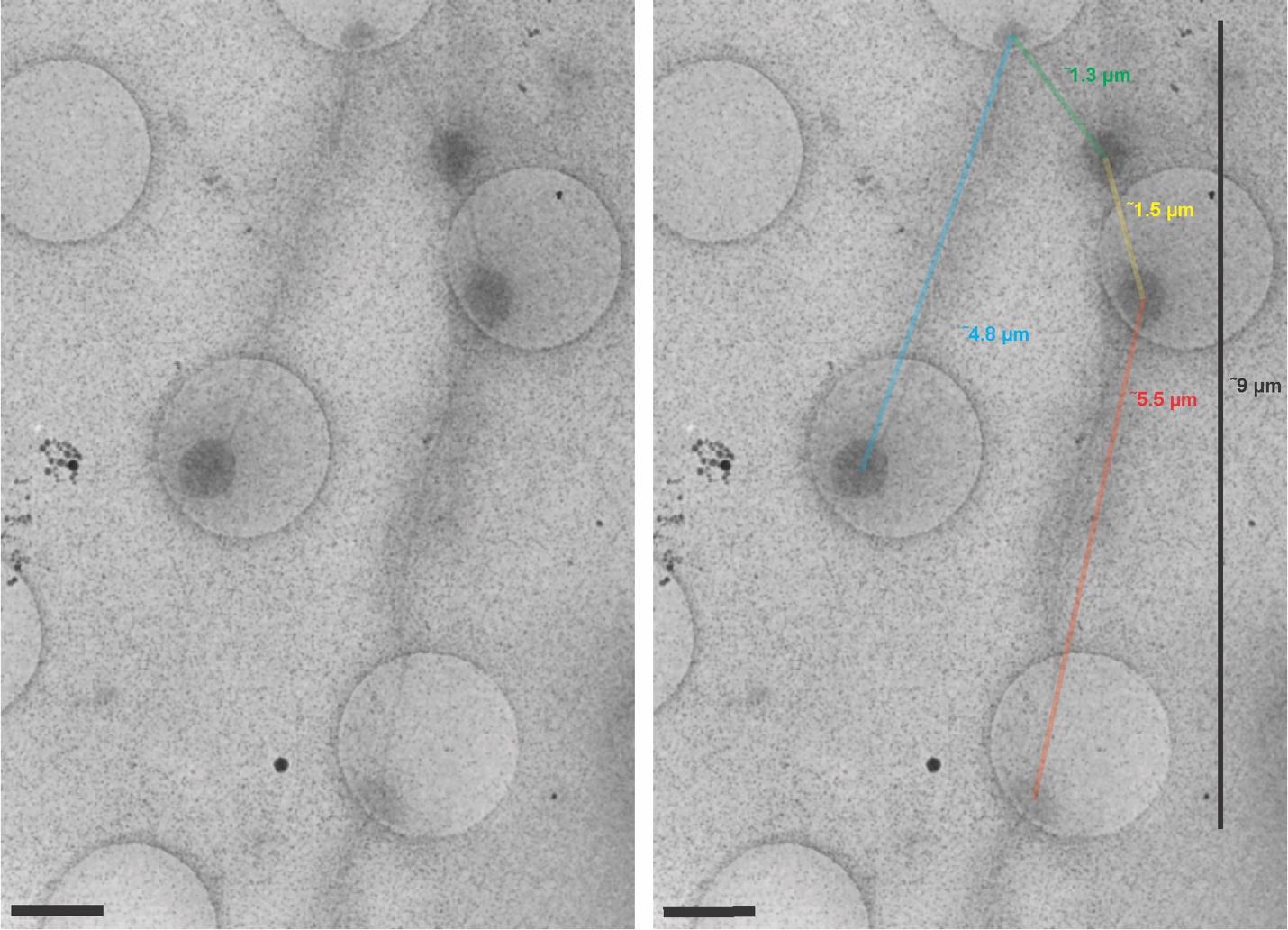

**Fig. S1. Low-magnification 2-dimensional (2D) projection images of Loki-ASV2 enrichment cultures derived from microbial mats.** Images revealed sparsely distributed round cell bodies connected by long extensions. Scale 1 µm.

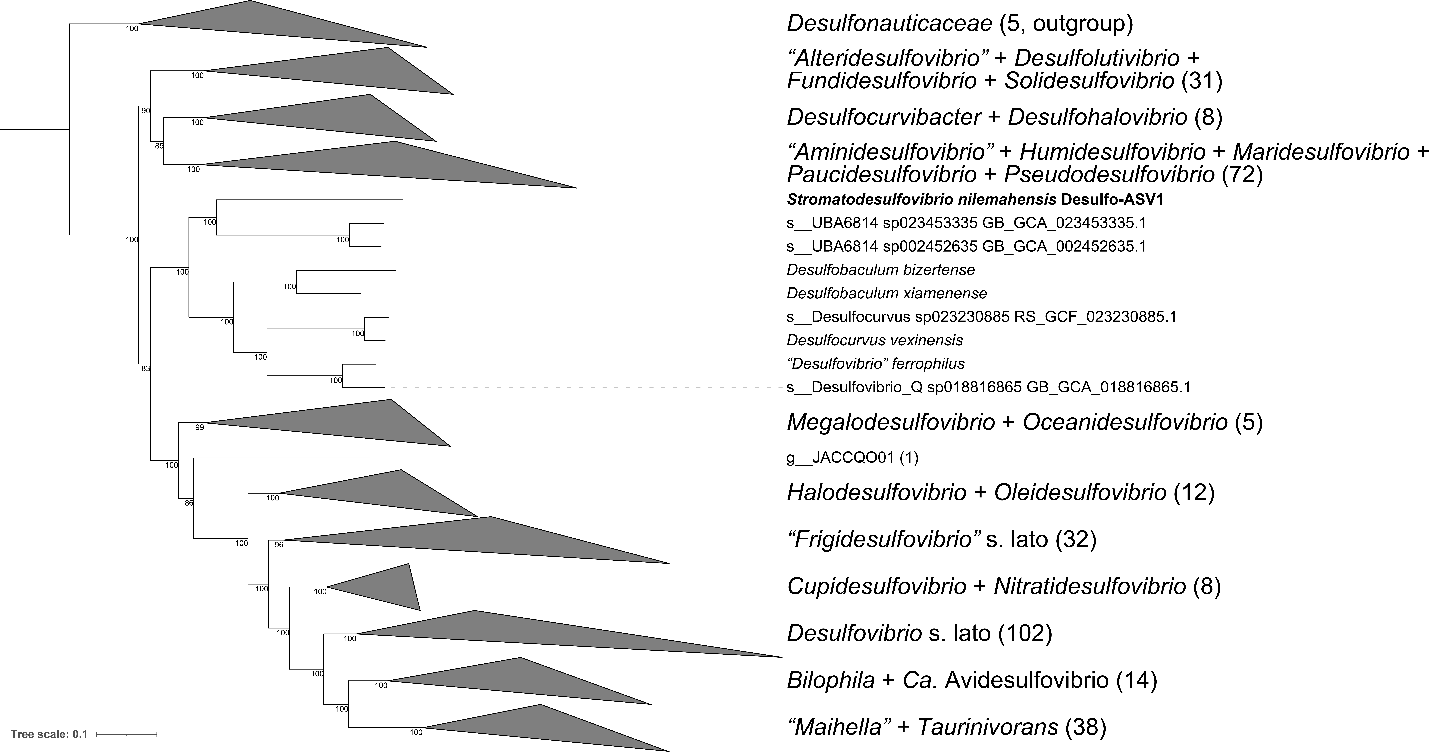

**Figure S2. *Desulfovibrionaceae* concatenated protein tree.** Tree was built using 115 single copy markers (Data S2) under the LG+F+I+R10 substitution model and 1,000 ultrafast bootstrap replicates. Collapsed nodes are identified by the names of the described genera contained within. Numbers at nodes indicate support values. *Desulfonauticaceae* was used as outgroup.

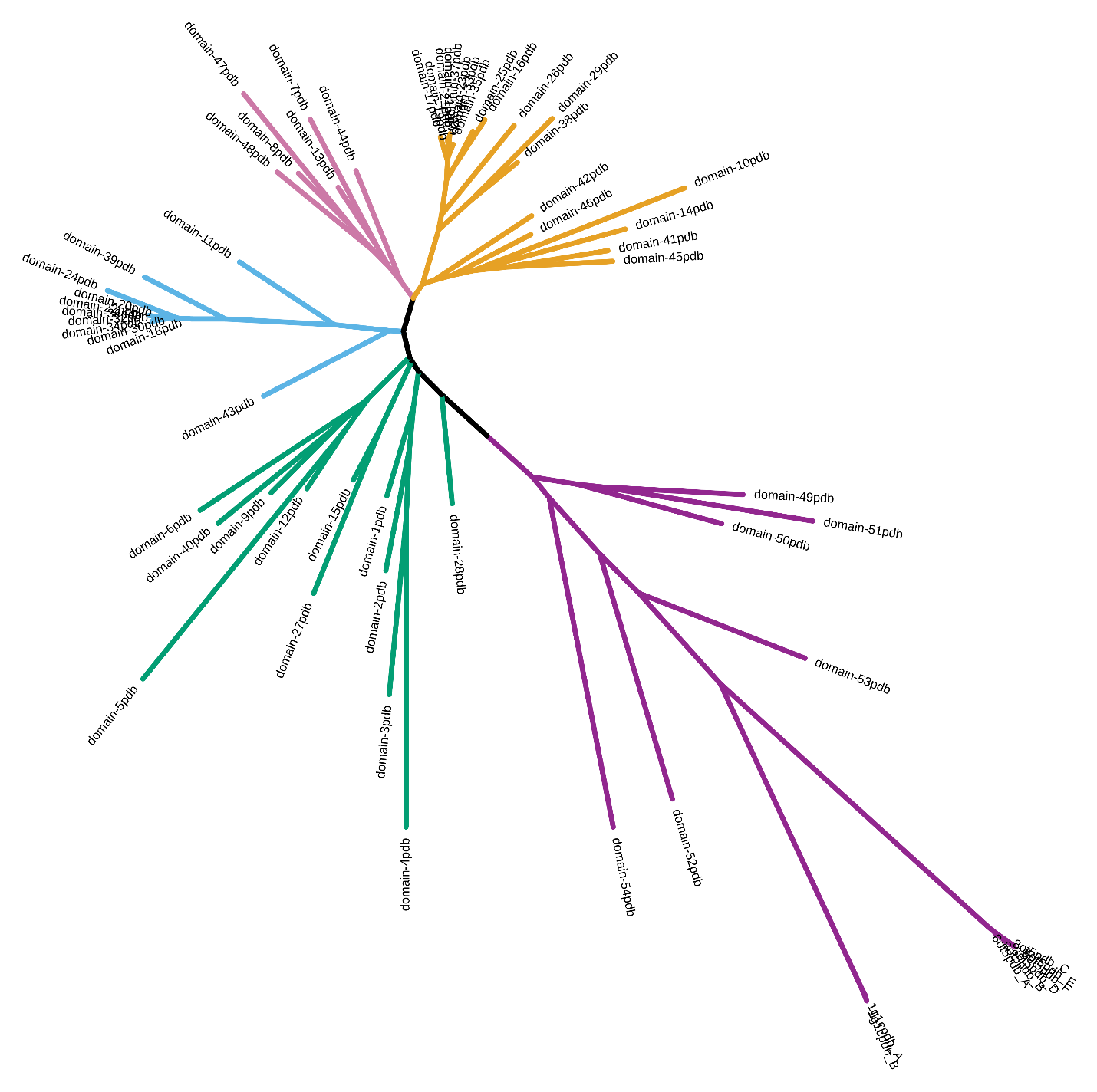

**Figure S3. Structural phylogeny of repeated fibronectin type-III-like domains in LOKIASV2_19760**. A maximum-likelihood phylogeny based on partitioning the amino acid and 3Di alignment from FoldMason. Domain subtypes were assigned and coloured as per Figure 3B.

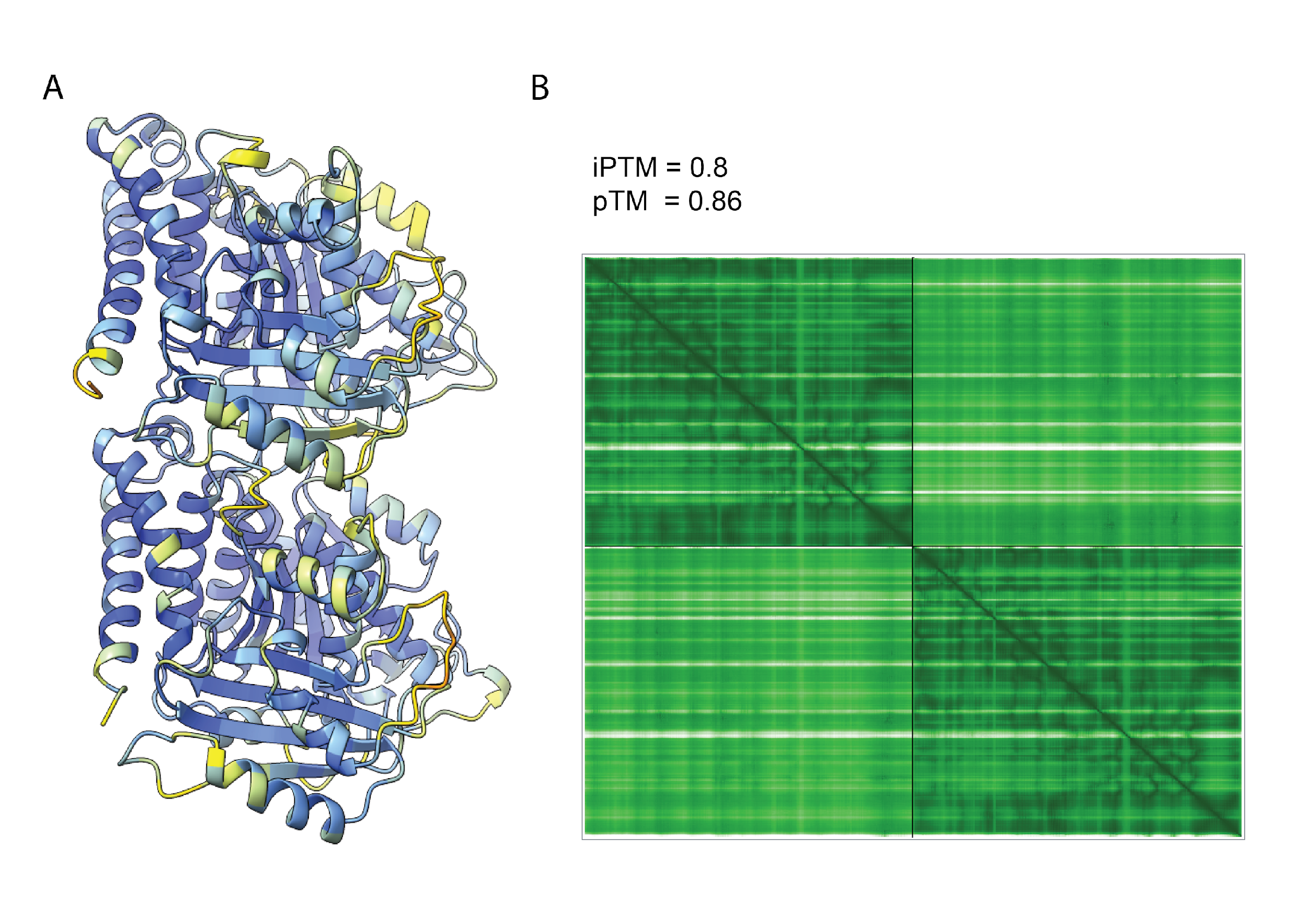

**Figure S4.** **AlphaFold3 model of LOKIASV2_32180 (AtubA) and LOKIASV2_32170 (AtubB).** (A) Heterodimer model colored by pLDDT using the ChimeraX AlphaFold palette. (B) Confidence scores of the relative positions (iPTM) and overall structure (pTM) and graph of the predicted alignment error (PAE).

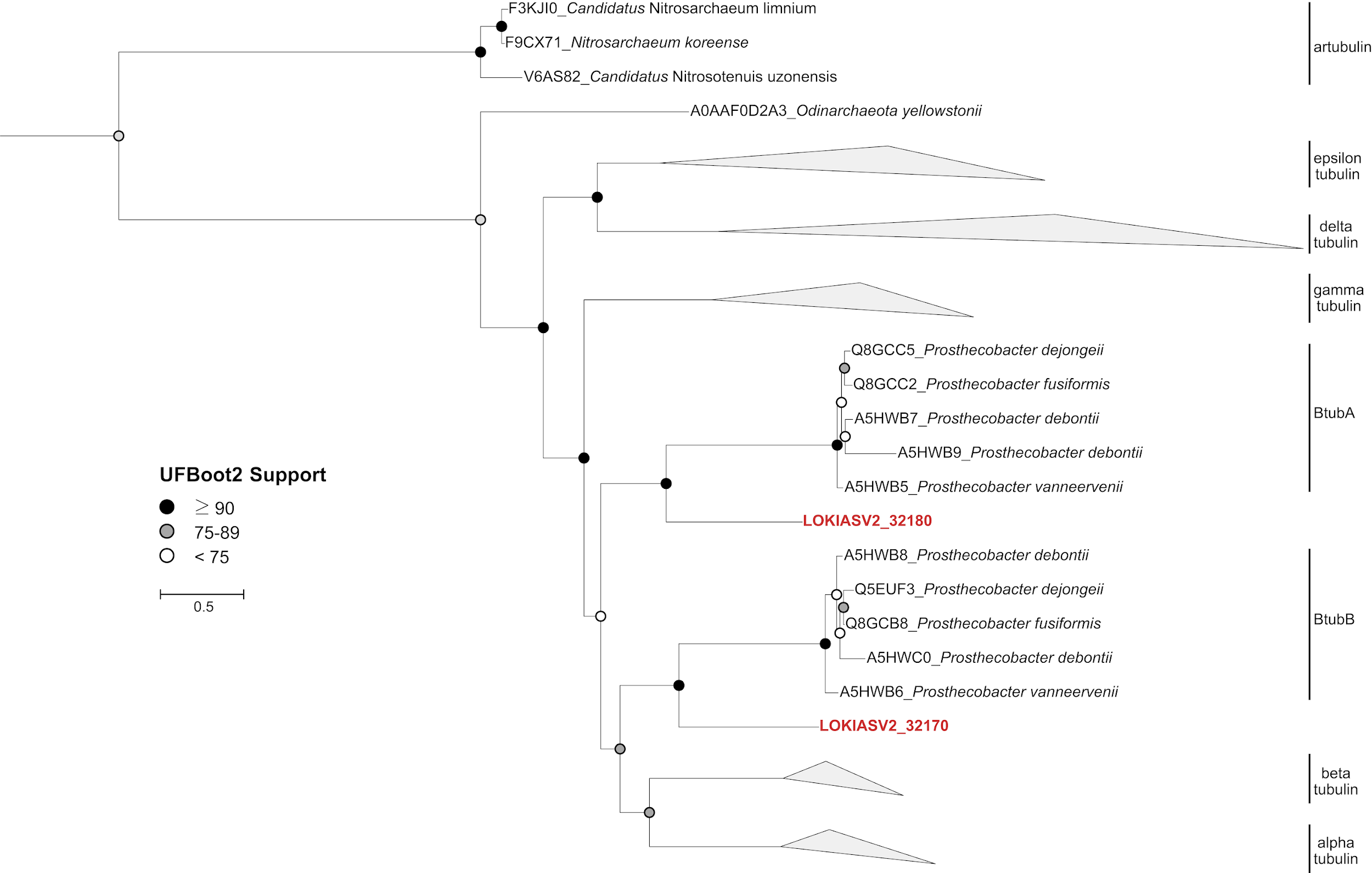

**Figure S5. Maximum-likelihood phylogeny of eukaryotic-like tubulins**. Two tubulins from *N. marumarumayae* Loki-ASV2 (red) were included in an alignment of sequences compiled in a previous study (*3*). Other nodes are labelled with the protein UniProt accession and originating organism. The tree was rooted to the archaeal artubulin group. To build the tree, the protein sequences were aligned using MAFFT-linsi (v7.481) and trimmed using ClipKit (v2.3.0) using the kpic-gappy option. The phylogeny was inferred with IQ-TREE (v2.2.2.5) using ModelFinderPlus and branch supports were calculated with 10,000 ultrafast bootstrap replicates.

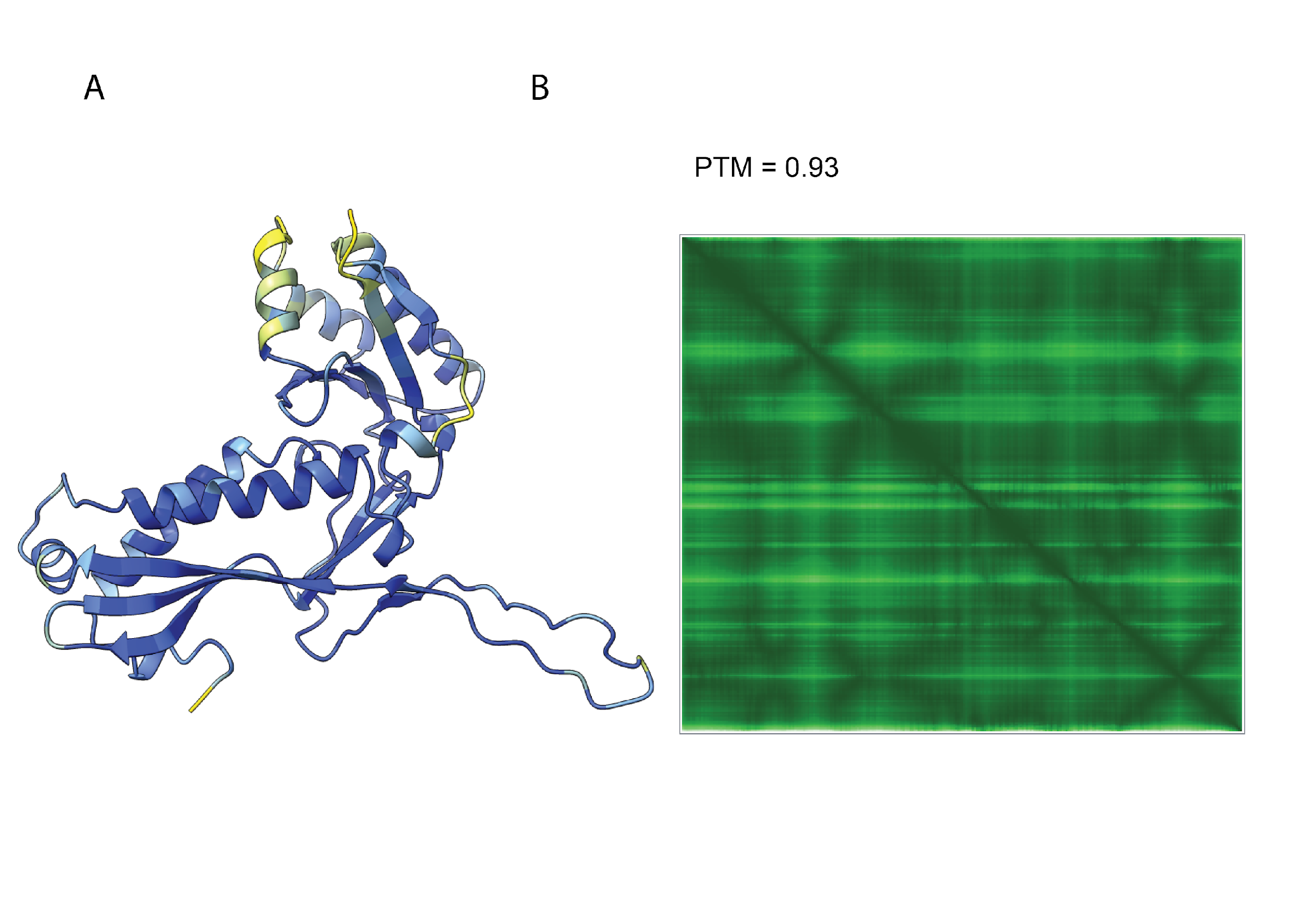

**Figure S6.** **AlphaFold3 model of LokiASV2_08240** (A) colored by pLDDT using the ChimeraX AlphaFold palette. (B) Confidence score of the overall structure (pTM) and graph of the predicted alignment error (PAE).

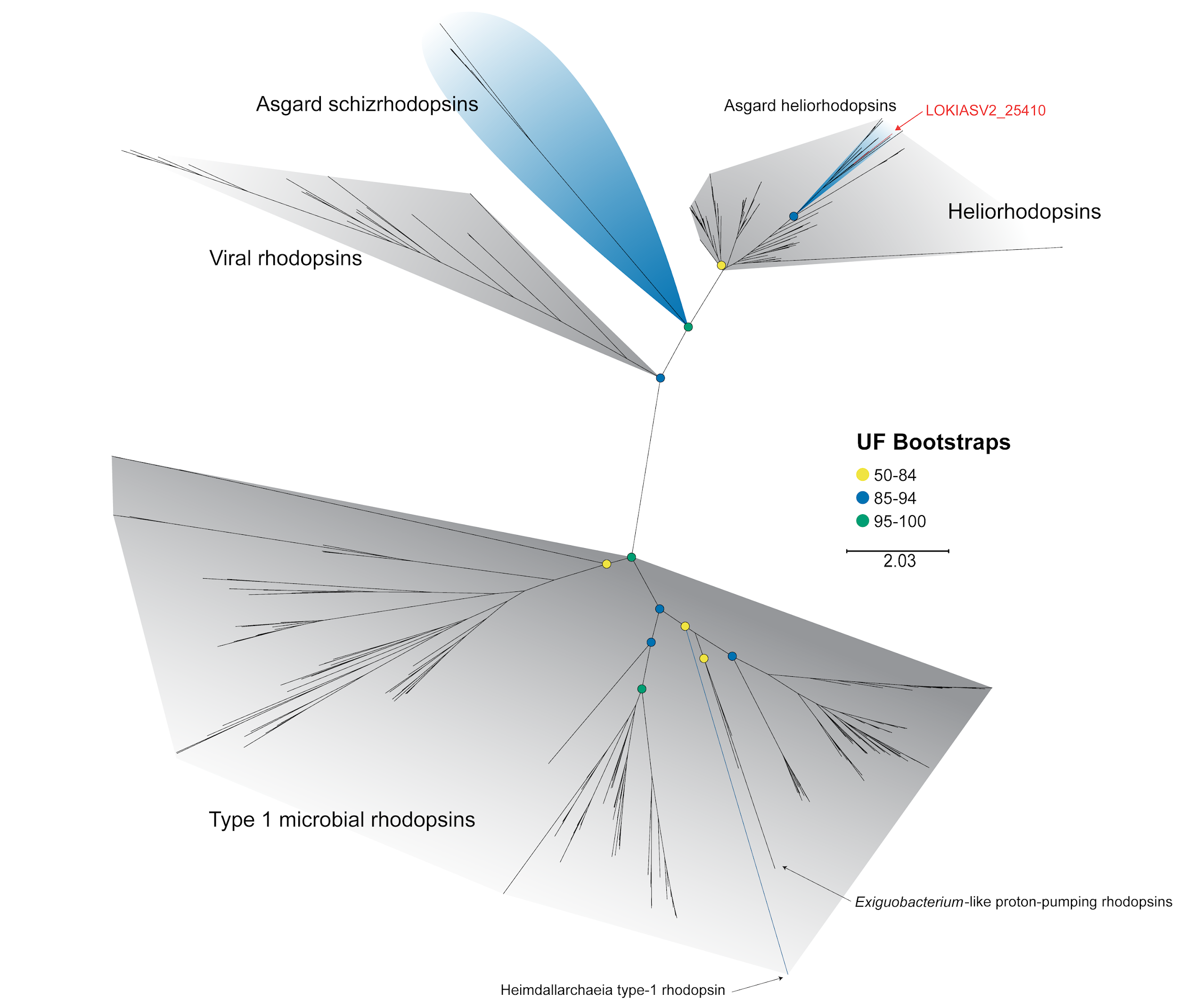

**Figure S7. Phylogeny of heliorhodopsin showing placement of LOKIASV2_25410**. A maximum-likelihood phylogenetic tree was created with IQ-TREE (v2.2.5) using ModelFinderPlus (*137*) and 10000 ultrafast bootstrap replicates.

**Supplementary Tables**

**Table S1.** Genome assembly details for *Nerearchaeum marumarumayae* Loki-ASV2 and the reference circularized *Promethearchaeaceae* MAGs.

| **Statistic** | **Loki-ASV2** ^a^ | **B35** ^b^ | **FW102** ^c^ | **MK-D1** ^d^ |
| --- | --- | --- | --- | --- |
| **Total Sequence Length (bp)** | 5,262,386 | 6,035,313 | 4,361,485 | 4,427,796 |
| **Completeness (actual)** ^e^ | 90.92 (100) | 91.26 (100) | 93.56 (100) | 92.65 (100) |
| **Contamination (actual)** ^e^ | 6.95 (0) | 9.15 (0) | 4.52 (0) | 6.15 (0) |
| **GC content (%)** | 34.0 | 33.6 | 31.0 | 31.2 |
| **Number of CDSs** | 4,544 | 5,120 | 3,695 | 3,946 |
| **CDSs in orthologs** ^f^ | 3,383 | 3,665 | 2,874 | 3,126 |
| **Average Protein Length** | 339.3 | 346.0 | 336.6 | 333.3 |
| **Coding Ratio (%)** | 87.9 | 88.1 | 85.5 | 89.1 |
| **Number of rRNAs (16S/23S/5S)** ^g^ | 2/2/2 | 3/3/3 | 2/2/3 | 1/1/1 |
| **Number of tRNAs (unique)** ^gh^ | 46 (22) | 46 (22) | 43 (22) | 46 (22) |
| **Number of CRISPRs** | 1 | 0 | 3 | 0 |

^a^: *Nerearchaeum marumarumayae* Loki-ASV2.

^b^: *Ca.* Lokiarchaeum ossiferum B35.

^c^: *Ca.* Harpocratesius repetitus FW102.

^d^: *Promethearchaeum syntrophicum* MK-D1.

^e^: Completeness and contamination estimates as per CheckM2.

^f^: CDSs with orthologs in other cMAGs in the table, based on analysis with Broccoli.

^g^: based on Rfam search with Infernal.

^h^: includes the 20 unique tRNA-aa plus tRNA-iMet and tRNA-SeC.

**Table S2.** Genome assembly details for *Stromatodesulfovibrio nilemahensis* Desulfo-ASV1.

| **Statistic** | **Desulfo-ASV1** ^a^ |
| --- | --- |
| **Total Sequence Length (bp)** | 4,134,807 |
| **Completeness (actual)** ^b^ | 97.06 (100) |
| **Contamination (actual)** ^b^ | 0.92 (0) |
| **GC content (%)** | 61.0 |
| **Number of CDSs** | 3,779 |
| **Average Protein Length** | 324.0 |
| **Coding Ratio (%)** | 88.8 |
| **Number of rRNAs (16S/23S/5S)** ^c^ | 2/2/2 |
| **Number of tRNAs (unique)** ^cd^ | 51 (22) |
| **Number of CRISPRs** | 0 |

^a^: *Stromatodesulfovibrio nilemahensis* Desulfo-ASV1.

^b^: Completeness and contamination estimates as per CheckM2.

^c^: based on Rfam search with Infernal.

^d^: includes tRNA-fMet and tRNA-SeC.

| **Table S3. Structural alignment statistics for Loki-ASV2 AtubA/B and mammalian alpha/beta tubulin.** | | | | | |
| --- | --- | --- | --- | --- | --- |
| Alignment type | Chain pairing observed | *Pruned atom pairs* | | *All atom pairs* | |
|  |  | No. pairs | RMSD (Å) | No. pairs | RMSD (Å) |
| AtubA *vs* α/β-tubulin (bb) | AtubA to β-tubulin | 358 | 1.091 | 419 | 1.950 |
| AtubB *vs* α-tubulin (bb) | AtubB to α-tubulin | 361 | 1.040 | 411 | 2.313 |
| AtubAB *vs* α/β-tubulin (ss) | AtubA to β-tubulin and AtubB to α-tubulin | 549 | 1.195 | 830 | 2.745 |

**Supplementary Movies and Data descriptions.**

**Movie S1. (separate file)**

3D tomogram reconstruction and segmentation of Loki-ASV2, an Asgard archaeon enriched from microbial mats. Scale bars 100 nm.

**Movie S2. (separate file)**

Interaction between Loki-ASV2 and Desulfo-ASV1 visualized by cryoET and segmentation. Scale bars 100 nm.

**Data S1. (separate file)**

Enrichment culture G2.24 16S rRNA gene amplicon relative abundance.

**Data S2. (separate file)**

Data associated with the phylogenomic and comparative genomic analyses of Loki-ASV2 and Desulfo-ASV1. Includes list of genomes and markers included in phylogenomic analyses, substitution models used in the individual/partitioned trees, average amino acid identity with closest relatives, and a summary of syntenic blocks shared between Loki-ASV2 and reference circularized MAGs.

**Data S3. (separate file)**

A. Full uncollapsed *Promethearchaeales* concatenated protein tree. Tree was built using 47 single copy markers (Data S2) under the LG+F+R8 substitution model and 1,000 ultrafast bootstrap replicates. Circles at internal nodes indicate ultrafast bootstrap support values. Order o__Helarchaeales was used as outgroup.
B. Partitioned maximum likelihood phylogenetic tree of *Promethearchaeales*. Tree was built using 47 single copy markers and 1,000 ultrafast bootstrap replicates. Partitions and their substitution models were defined based on the constituent markers (Data S2). Circles at internal nodes indicate ultrafast bootstrap support values. Order o__Helarchaeales was used as outgroup.
C. *Promethearchaeales* phylogenetic tree under the multispecies coalescence model (supertree). Tree was built with the weighted ASTRAL (hybrid) algorithm using 47 individual single marker ML trees as input (Data S2). Circles at internal nodes indicate support values. Order o__Helarchaeales was used as outgroup.

**Data S4. (separate file)**

A. Full uncollapsed *Desulfovibrionaceae* concatenated protein tree. Tree was built using 115 single copy markers (Data S2) under the LG+F+I+R10 substitution model and 1,000 ultrafast bootstrap replicates. Circles at internal nodes indicate ultrafast bootstrap support values. *Desulfonauticaceae* was used as outgroup.
B. Partitioned maximum likelihood phylogenetic tree of *Desulfovibrionaceae*. Tree was built using 115 single copy markers and 1,000 ultrafast bootstrap replicates. Partitions and their substitution models were defined based on the constituent markers (Data S2). Circles at internal nodes indicate ultrafast bootstrap support values. *Desulfonauticaceae* was used as outgroup.
C. *Desulfovibrionaceae* phylogenetic tree under the multispecies coalescence model (supertree). Tree was built with the weighted ASTRAL (hybrid) algorithm using 115 individual single marker ML trees as input (Data S2). Circles at internal nodes indicate support values. *Desulfonauticaceae* was used as outgroup.

**Data S5. (separate file)**

Genes encoding metabolic proteins and associated pathways in Loki-ASV2 and Desulfo-ASV1 genomes.

**Data S6. (separate file)**

Strategies to enrich for Asgard archaea that were tested and optimized over the course of this study.

**Data S7. (separate file)**

List of cytoskeletal and morphological proteins encoded in Loki-ASV2 genome.

**Data S8. (separate file)**

Curated set of Loki-ASV2 pdb files used as queries against the custom database and searched using foldseek and reverse foldseek.
