## Supplementary figures and images for "An Asgard archaeon from a modern analog of ancient microbial mats"

### Data S3

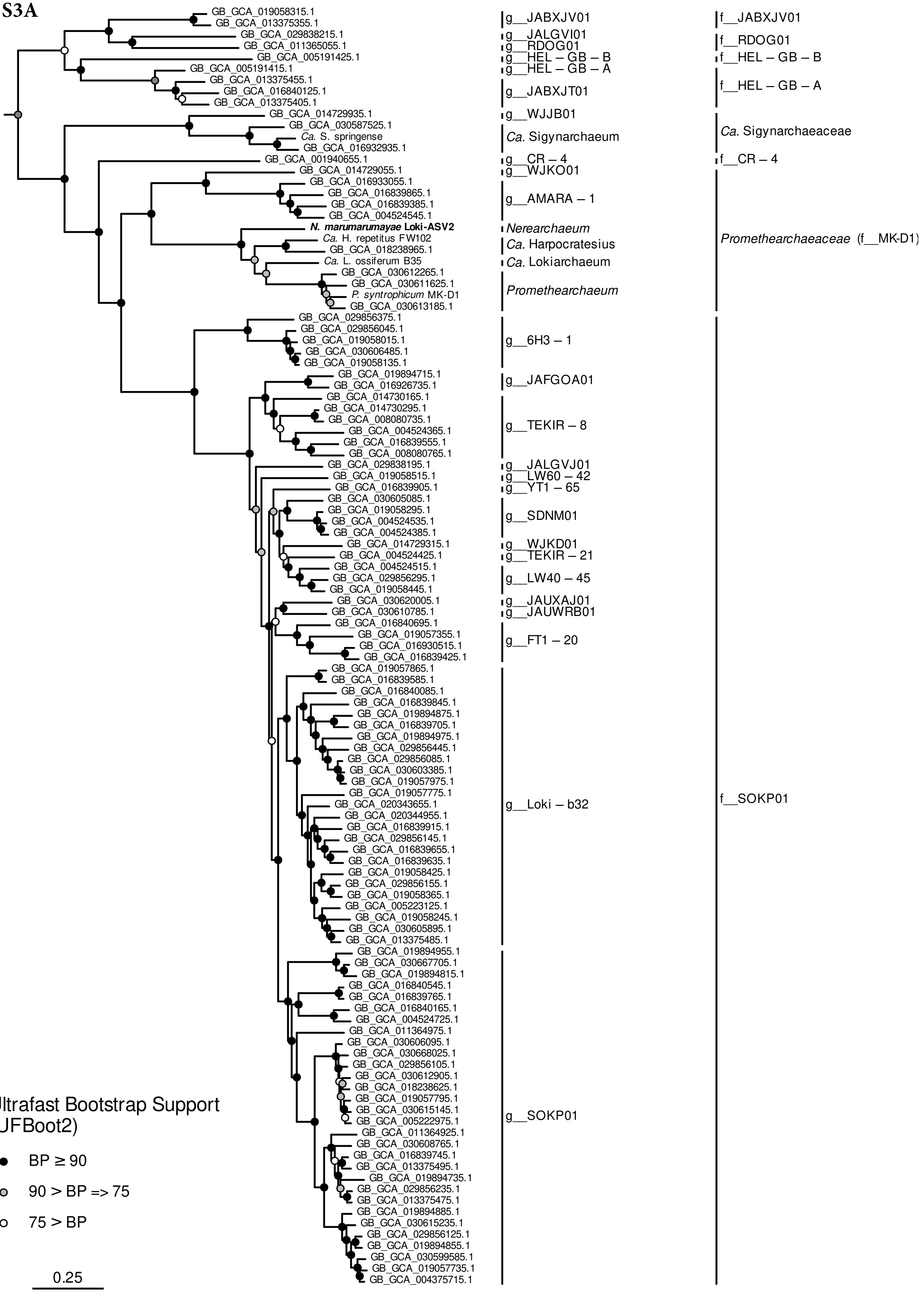

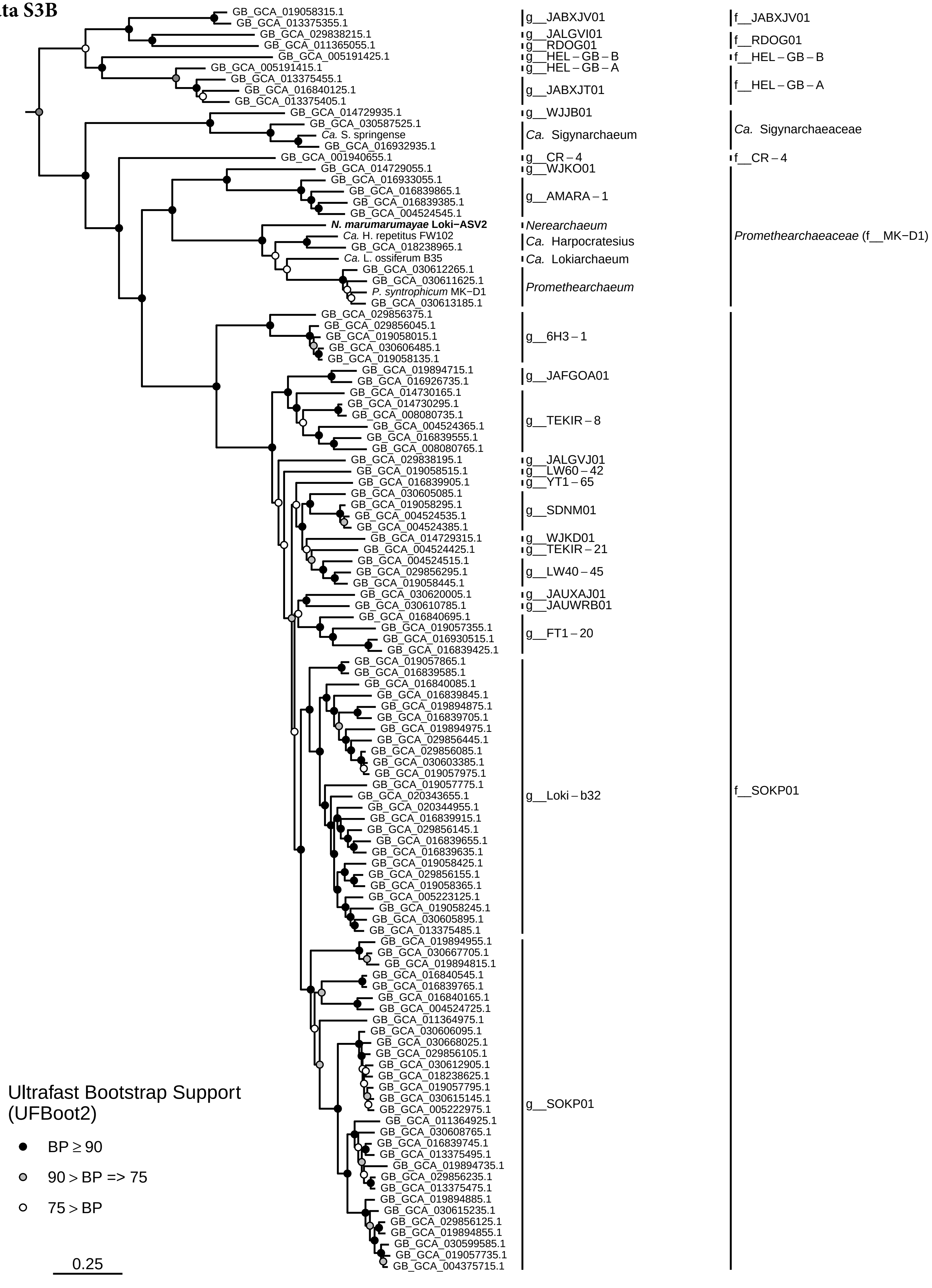

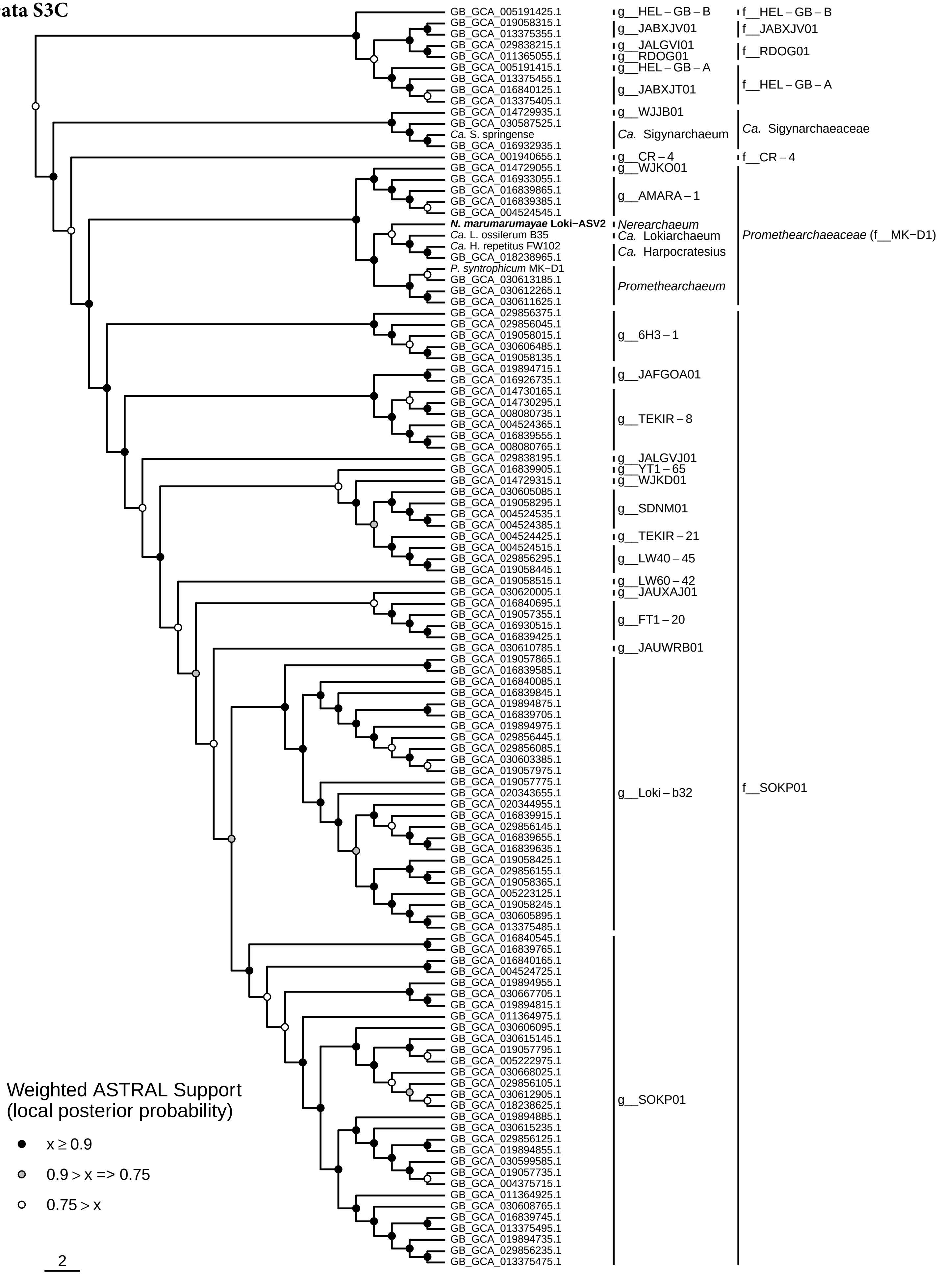

### Data S4

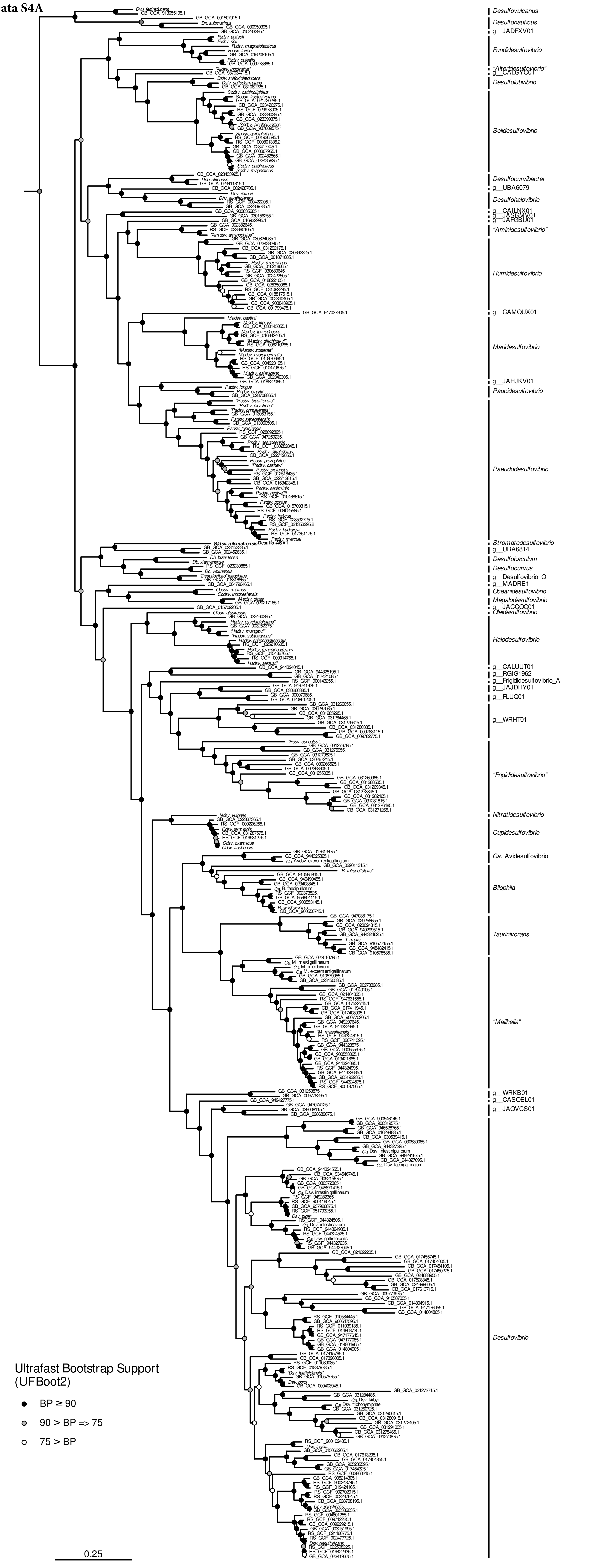

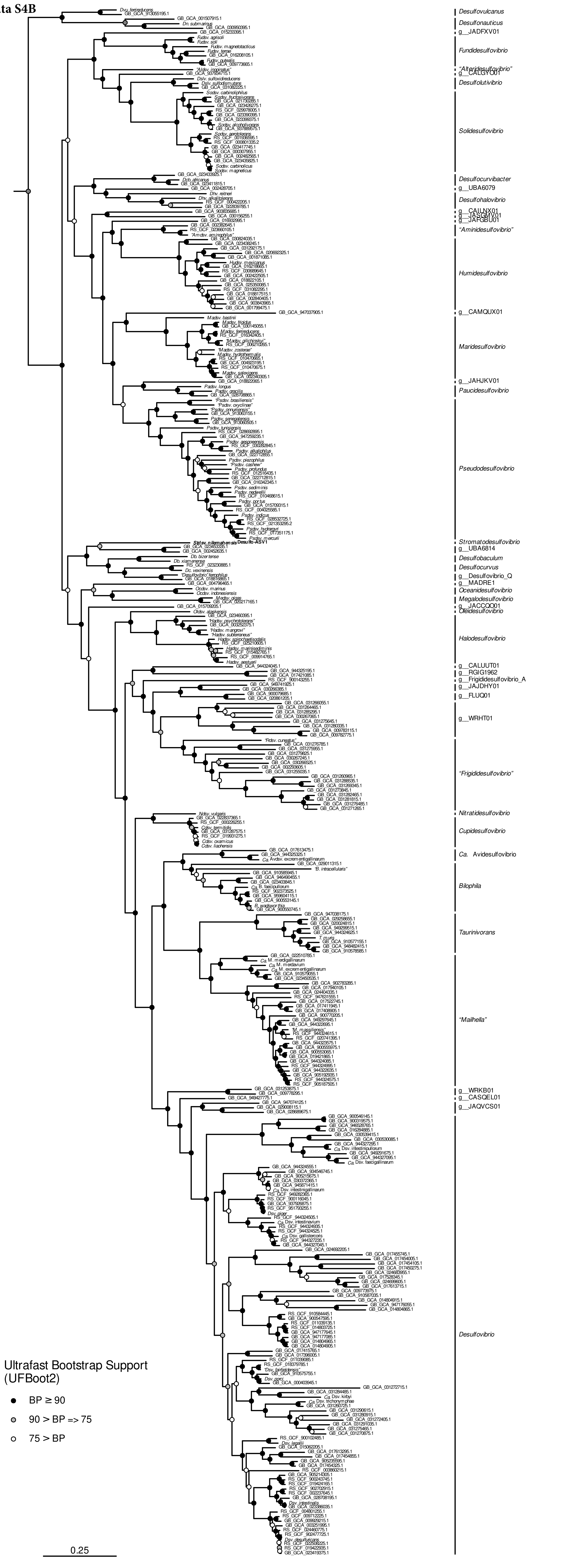

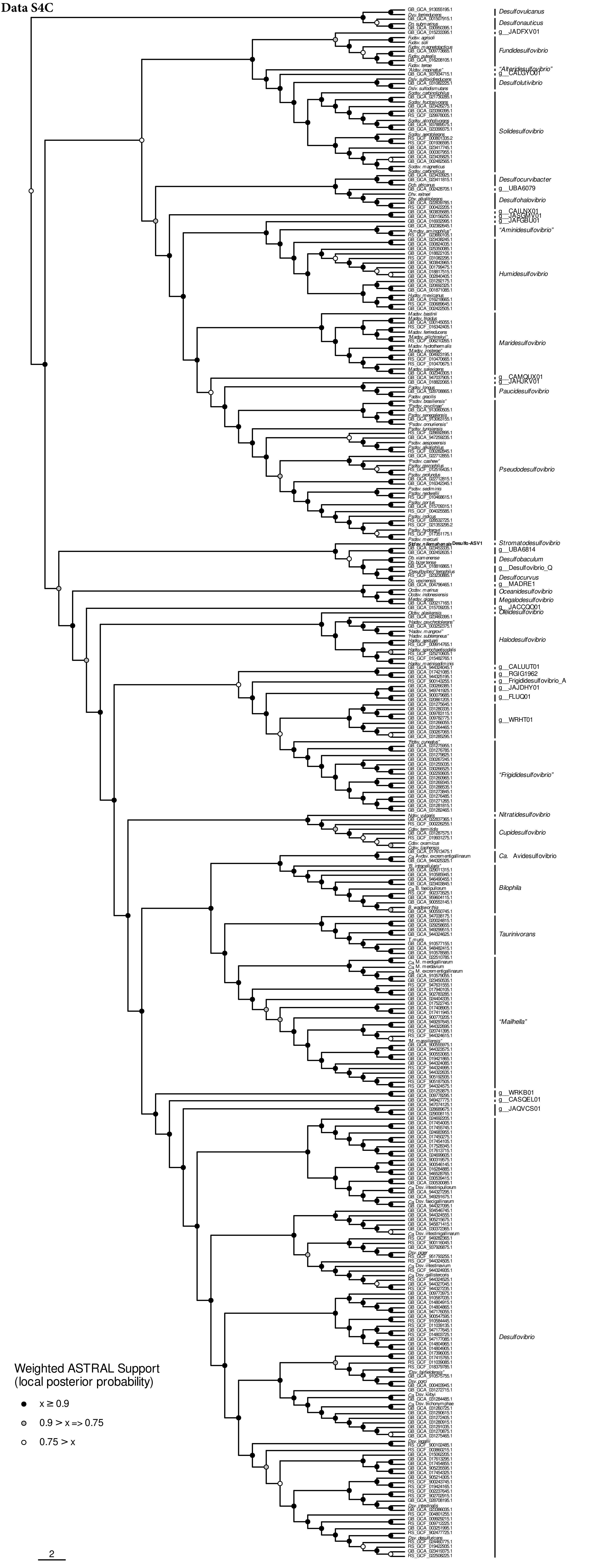
